# Optimal Predictivity of Random Signatures: Causes and Good Modeling Practices

**DOI:** 10.64898/2026.11.13.764046

**Authors:** Sisi Ma, Roshan Tourani, Linmin Wang, Mehmet Ugurbil, Thomas Kirsh, Jinhua Wang, Boris Winterhoff, Alexander Statnikov, Constantin Aliferis

**Affiliations:** University of Minnesota; New York University

## Abstract

In high-throughput omics data, randomly-selected sets of markers (“signatures”) have large and often optimal phenotypic predictivity. This creates problems of over-interpretation of biological and predictive importance of signatures, of bioinformatics algorithms and expert knowledge-driven selection.

We conduct first an empirical study establishing that the phenomenon occurs across a variety of types of datasets/omics types and algorithms (224 combinations, >120 million models fitted). Random signatures are systematically reproducible across datasets. We develop a comprehensive theoretical framework, encompassing definitions, feasibility and impossibility theorems, and analytical formulas. The theory reveals as necessary and sufficient causes of these anomalous behaviors, the existence of large equivalence classes of optimally predictive feature sets, and their structure of information equivalences.

We validate the theory with additional experiments and propose good practices intended to strengthen scientific rigor. Our findings apply to both algorithmic discovery and expert knowledge-driven signature selection, and to both existing and future algorithms and expert criteria. They also extend to all types of data with similar equivalence classes of optimal predictors.

## Introduction

Discovery of molecular signatures is a staple of modern biomedical research with numerous basic science and translational and clinical science uses^1–8^, with >210,000 articles tagged in Pubmed with a “molecular signature” keyword. Supplementary Document 2 discusses examples from cellular senescence and cancer motivating the present research. One major difficulty in discovering reproducible and biologically relevant markers^a^ and signatures^b^ from high-dimensional omics data, is that the signatures identified by state-of-the-art computational algorithms are strongly predictive of phenotypes of interest and as originally reported^13^ twenty-eight of studied breast cancer signatures (60%) were not significantly better outcome predictors than random signatures of identical size. This creates three problems: (a) Over-interpretation of signature biological importance on account of strong predictivity (i.e., despite that random signatures of the same size have same predictivity with high probability). (b) Under- and over-interpretation of predictive importance: under-interpreting when signatures are rejected because they are not statistically significantly different than random, yet they are reproducible and optimally accurate; over-interpreting when the measured high predictivity is viewed as possibly unique, although shared by many (unexamined) random signatures. (c) Over-interpretation of the merits of bioinformatics algorithms or expert knowledge-driven selection: whereas the effort invested in developing and applying sophisticated algorithms or expert appraisal of the literature is wasted or irrelevant, if they do not add significantly to random selection.

Compounding the above, a staple of biomedical research is to test findings against the null hypothesis^14^, which in our setting is that a discovered signature is not better than random. New markers and signatures reported in the omics literature, however, are typically not tested for being statistically significantly different than random, even among the studies with the highest impact^15,16^. New or existing algorithms^17^ as well as human expert selection^18^ are also typically not tested for added value over random selection. Some authors adopt the assumption that biologically meaningful markers/signatures reproduce better than randomly selected biomarkers/signatures; and, if a marker set is better than random performance, it is biologically meaningful^19,20^.

In the present report, we provide a comprehensive analysis of these issues: (a) we first show empirically that the extent of the randomness problem is ubiquitous, in terms of affected omics molecules/assay types, phenotypes and algorithms used, far beyond the omics types, algorithms, and research fields originally reported^13^. We then provide a complete theoretical account of the phenomenon and we test and validate the theory with extensive experiments. (b) Our analysis reveals that the root cause of the phenomenon is the existence of a *large set of phenotype information-equivalent and predictively-optimal marker sets (aka “Markov Blankets”) in omics data*. The effect of this equivalence class is in turn determined by (i) its size and (ii) the information equivalence structure of the embedded equivalence class of *predictively optimal and non-reducible marker sets (aka “Markov Boundaries”)*. These equivalence classes provide the necessary and sufficient conditions for random biomarkers and signatures performing predictively optimally, described by precise mathematical formulas. (c) We examine the *structural and statistical* reasons for the equivalence classes, and demonstrate their existence in real-life datasets. (d) We differentiate between predictive vs causal (i.e., mechanistic) discovery and show that several widely-used algorithms do not have benefits over random biomarker discovery. Testing for randomness does not inform the mechanistic significance of markers and signatures in TIE distributions. Alternative reliable strategies for investigating mechanistic significance are discussed. We finally (e) codify a number of proposed good modeling and interpretation practices regarding marker/signature randomness aiming to help protect biological and translational investigators from related inadequate modeling or misinterpretation of results.

## Results

### Empirical behavior of algorithms and random signature selection across various omics data

#### Random feature selection can result in excellent and generalizable predictive performance

These results hold for various types of omics assays (not just transcriptomics) and for both bulk and single cell data. The predictivity increases as the number of features increases across 14 omics datasets examined in the current study. In general, the rate of AUC increase is more rapid at smaller marker set sizes, but slower as the feature set size grows larger (Figure 1a). The performance of randomly selected features generalizes to external datasets (Figure 1b).

**Figure 1.**
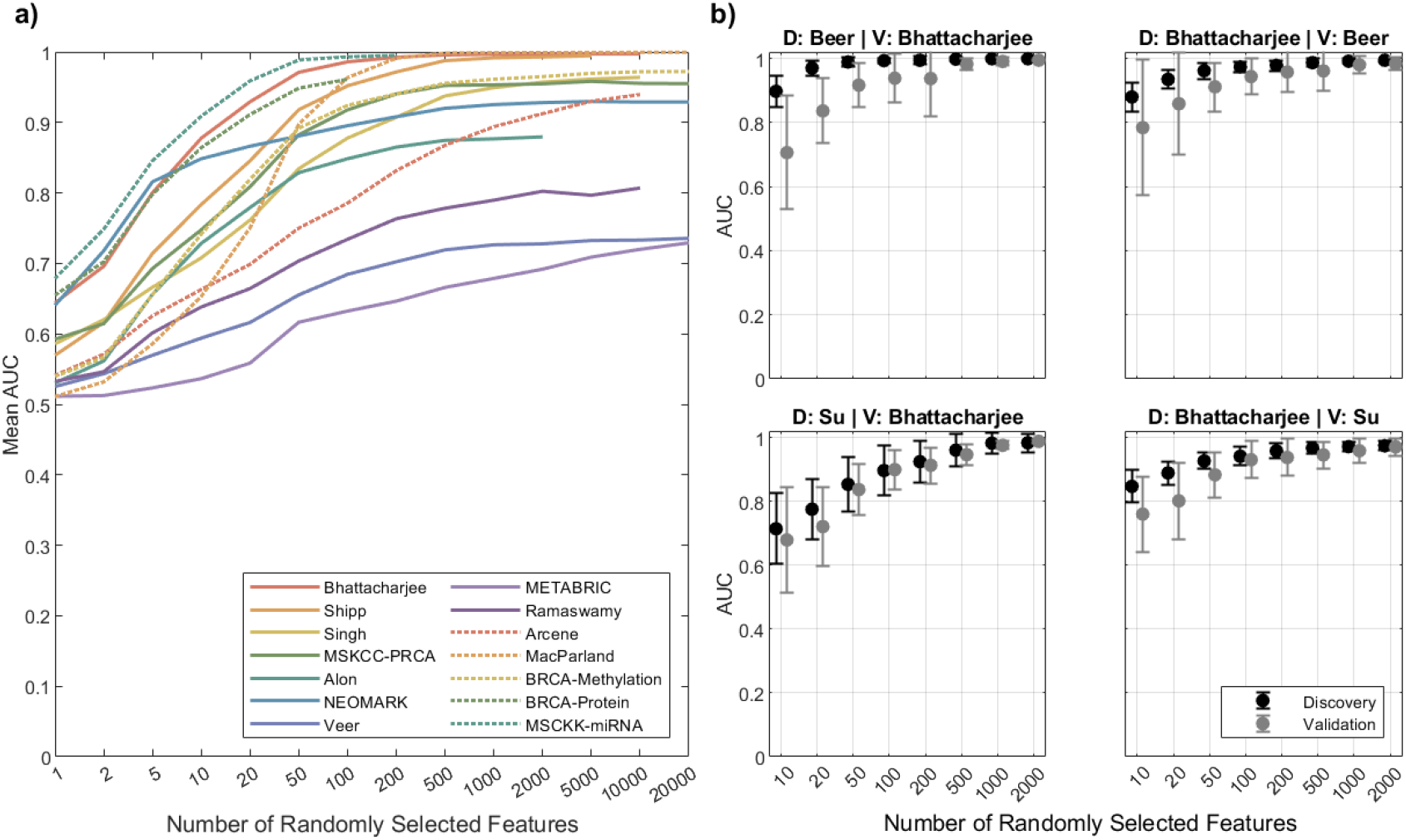
Empirical results of random feature selection on omics data. (a) As the number of randomly selected features increases, the predictivity of random feature selection increases. Solid lines represent gene expression datasets, dotted lines represent other omics datasets. (b) The performance of random selected features on discovery datasets, reproduces to external validation datasets derived from independent studies. The performance variance between discovery and validation datasets decrease as the number of randomly selected features increases. More information is given on datasets and analysis procedures in Online Methods.

**Random feature selection will often achieve similar performance compared to algorithmic feature selection at matching feature set sizes. Some feature selection algorithms do not offer added value compared to random feature selection on average over all datasets.**

**Table 1** summarizes the performance of 8 state-of-the-art widely-adopted algorithms vs. random feature selection of the same feature set size (Online Methods). The performance of both algorithmic feature selection and random feature selection spans a rather wide range over the 14 omics datasets examined. The comparative performance of random feature selection vs. algorithmic feature selection is affected by the data distribution, the algorithm (its predictivity and number of features selected), and the variance of the estimator used for predictivity estimation.

**Table 1.** Empirical Results of algorithmic signature selection vs. random signature selection in omics data. Predictive performance of 8 algorithmic feature selection methods in 14 omics datasets using two different performance estimators, the repeated cross validation predictivity estimator (RPE) and the non-repeated cross-validation estimator (NRPE). AUC(FS): the AUC of the algorithmic feature selection method; AUC(RFS): the AUC of the random feature selection with size of signatures equal to the algorithmic method’s output for that dataset per cross validation fold (Online Methods); AUC(FS)-AUC(RFS): added value of the algorithmic feature selection over random feature selection. Better than random: number of datasets where the algorithms outperform random selection. Cells contain the mean and range ([min, max]) of the reported metrics over the 14 datasets (averaged over cross validation folds in each dataset). Bold font indicates that performance is statistically indistinguishable from the best performance in the same row/metric (FDR q-value threshold=0.05).

| Estimator |  | GLL-OPT | RFVS | LARSEN | UAF-KW1 | SIMCA | SVM-RFE | RELIEF-SVM | T-fdr |
|---|---|---|---|---|---|---|---|---|---|
| RPE | **AUC(FS)** | **0.91 [0.72,1.00]** | 0.86 [0.66,1.00] | 0.89 [0.69,1.00] | 0.88 [0.69,1.00] | 0.85 [0.54,1.00] | 0.89 [0.71,1.00] | 0.84 [0.65,1.00] | **0.88 [0.59,1.00]** |
|  | **AUC(RFS)** | 0.74 [0.58,0.94] | 0.68 [0.54,0.95] | 0.73 [0.61,0.85] | 0.69 [0.60,0.85] | 0.73 [0.54,1.00] | 0.70 [0.60,0.83] | 0.75 [0.63,0.85] | **0.84 [0.52,1.00]** |
|  | **AUC(FS)-AUC(RFS)** | **0.17 [0.02,0.33]** | **0.18 [0.05,0.31]** | **0.16 [0.02,0.39]** | **0.19 [-0.08,0.40]** | 0.12 [-0.05,0.32] | **0.19 [-0.02,0.38]** | 0.09 [-0.06,0.28] | 0.04 [0.00,0.22] |
|  | **Number of Features** | **32 [3,220]** | **16 [2,109]** | 51 [2,280] | 1230 [1,13715] | 992 [2,7367] | 1675 [1,19850] | 1796 [3,18593] | 1920 [2,6725] |
|  | **Better Than Random** | 14/14 | 14/14 | 13/14 | 13/14 | 13/14 | 13/14 | 11/14 | 10/14 |
| NRPE | **AUC(FS)** | **0.91 [0.72,1.00]** | **0.88 [0.56,1.00]** | **0.91 [0.68,1.00]** | **0.90 [0.66,1.00]** | 0.84 [0.49,1.00] | 0.87 [0.63,1.00] | 0.84 [0.54,1.00] | **0.88 [0.59,1.00]** |
|  | **AUC(RFS)** | 0.75 [0.58,1.00] | 0.69 [0.54,0.96] | 0.74 [0.60,0.89] | 0.69 [0.58,0.85] | 0.73 [0.56,1.00] | 0.70 [0.52,0.87] | 0.74 [0.56,0.86] | **0.84 [0.52,1.00]** |
|  | **AUC(FS)-AUC(RFS)** | **0.16 [0.00,0.31]** | **0.19 [-0.00,0.31]** | **0.17 [0.02,0.38]** | **0.21 [-0.06,0.40]** | 0.12 [-0.10,0.30] | 0.17 [-0.03,0.38] | 0.09 [-0.10,0.30] | 0.04 [0.00,0.26] |
|  | **Number of Features** | 96 [3,906] | **17 [2,113]** | 51 [2,344] | 1672 [1,16654] | 1040 [2,7675] | 1403 [1,14266] | 2350 [2,26012] | 1937 [1,6727] |
|  | **Better Than Random** | 11/14 | 12/14 | 12/14 | 12/14 | 9/14 | 9/14 | 7/14 | 3/14 |

Notably, random feature selection can achieve excellent performance (e.g. AUC>0.9; Table 1, row 2, Supplementary Table S5). When using the 50x repeated (lower-variance) nested cross validation (NCV) predictivity estimator^21^, only two algorithmic feature selection methods, GLL-OPT and RFVS, statistically significantly outperform their corresponding random feature selection in all 14 omics datasets examined. These feature selection methods are also the methods that selected, on average, the smallest feature sets. The T-fdr and RELIEF-SVM outperformed random feature selection in 10 and 11 out of 14 datasets when using the RPE. These algorithms least frequently outperform random feature selection in our tests.

When using the non-repeated (higher variance) nested cross validation predictivity estimator (NRPE), all algorithms outperformed their corresponding random feature selection in a much smaller number of datasets (compare Table 1, row 5 vs. row 10),. Specifically, T-fdr and RELIEF-SVM in only 3 and 7 of 14 datasets respectively.

The added value of the algorithmic feature selection over the corresponding random feature selection ranges from 0.04 to 0.19 AUC averaged over all 14 datasets under the RPE. Out of the feature selection methods that have maximal statistically indistinguishable added value (Table 1, row 3, bolded cells), the GLL-OPT feature selection resulted in statistically significantly better performance compared to other algorithms (Table 1, row 1). T-fdr has statistically indistinguishable predictive performance compared to GLL-OPT, but compared to randomly selecting features of the same size, its added value over random is on average only 0.04 AUC points.

Overall in 37 out of 112 (33%) of datasets x algorithms with NRPE error estimation, algorithms did not outperform random selection of the same size. When using RPE, this happened in 8 out 112 (7%) analyses only, showing the importance of incorporating low-variance estimator protocols in omics analyses to reduce random selection effect^21^.

### Theoretical analysis

To systematically and quantitatively explain the above empirical results, and to facilitate the design and analysis of future studies, we developed a theoretical framework comprising 5 main Theorems and related definitions, and corollaries that together characterize the behavior of random feature selection. In the main text, we provide a summary description in language accessible to general science audiences. Formal definitions and proofs of the theorems geared to more technical audiences are provided in the **Supplementary Document 1**. We start with essential computational concepts employed in the analysis.

#### Essential concepts: Markov blankets, Markov boundaries, and how they are germane to signature optimal predictive performance

A Markov Blanket of a target phenotypic variable T (denoted by: MBL(T)) is a variable set that contains maximum information about T (i.e. we can use it to fit models from data with optimal predictive performance for T (Supplementary Document 1, Definition 2, Remark 2). A Markov Boundary (MB(T)) is a MBL(T) that is also minimal (aka non-reducible), i.e., removing any subset of variables from it will result in loss of predictive performance (Supplementary Document 1, Definition 1, Remark 1). Therefore, to achieve optimal performance for predicting T, a signature must be a MBL(T). If in addition, we need the signature to be most parsimonious, it must be exactly a MB(T). A non-minimal but optimally predictive signature (i.e., MBL(T)) may contain multiple MB(T)s and/or other unrelated variables.

In so-called “Faithful distributions” (a family that contains the vast majority of all theoretical distributions^22^, there is a unique MB^23^, E.g., Figure 2 bottom left panel illustrates a variable set with a unique MB(T)={A,B}. This set and any set that contains both A and B, contain maximal information for T, and thus can be used to fit a classifier resulting in optimal performance. There are 4 variable sets that can lead to optimal performance in the example. Removing A or B from the feature set reduces the performance for T. Variables X_1_ and X_2_ may contain information about T but do not add additional information to that of A and B.

**Figure 2.**
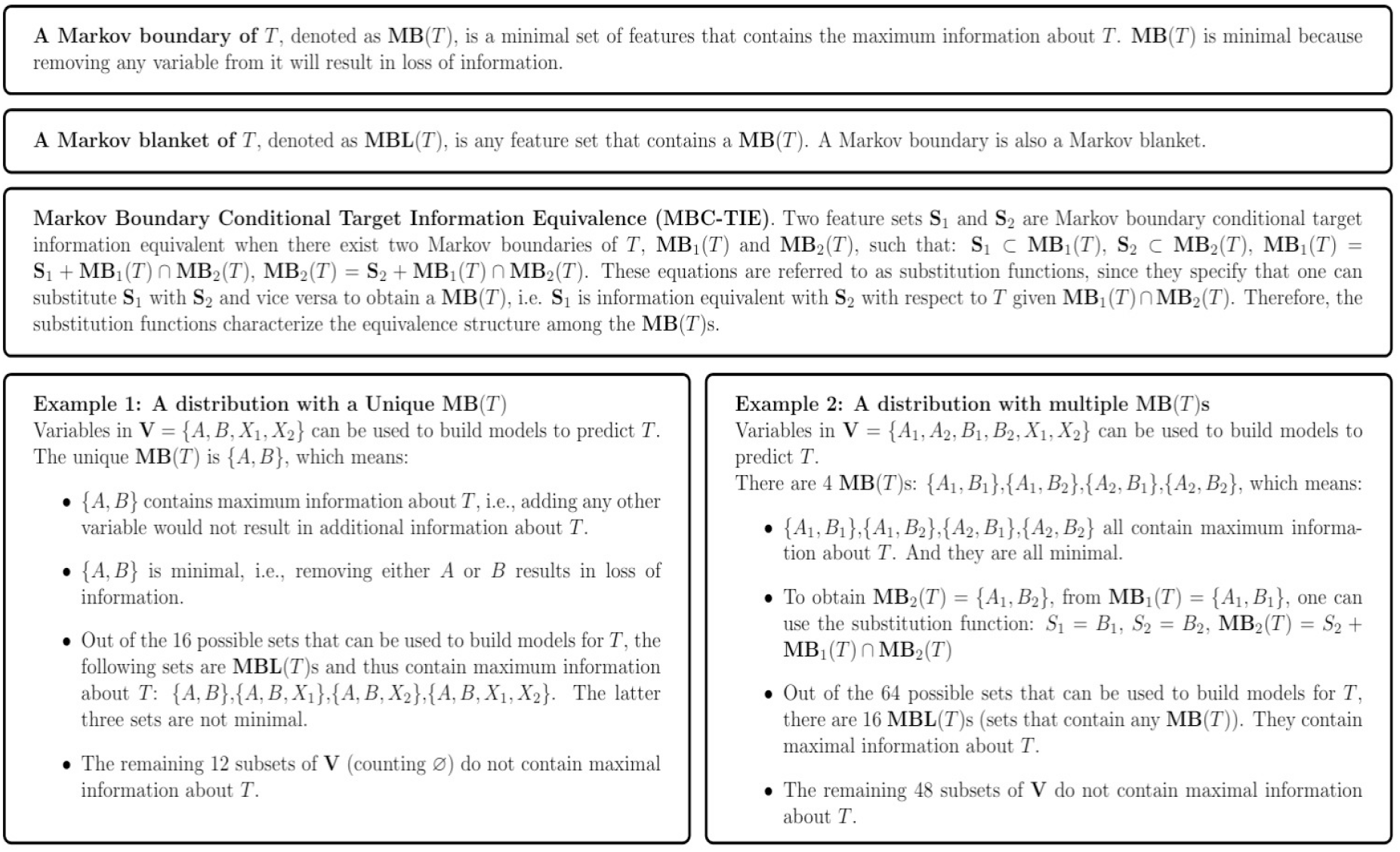
Markov boundaries & Markov blankets, optimal signatures and their equivalence classes.

In some non-faithful distributions, there are more than one MB(T), each one of them providing equivalent and maximal information about T^24^ (i.e. we call the phenomenon Markov Boundary Conditional Target Information Equivalence, MBC-TIE, Figure 2, Supplementary Document 1). Empirically, the presence of multiple MB(T) is often observed in biomedical data, especially in high-throughput omics data^8,24,25^. MBC-TIE can be created by a variety of mechanisms (information sharing among markers because of pathway biological connectivity, measurement assay engineered redundancy, or other structural reasons, plus statistical variance in small sample sizes leading to inability to differentiate the signal for T of some signatures).

The simple illustrative example in Figure 2 Example 2 shows a dataset with 4 MB(T)s. Any variable set containing A_1_ or A_2_, and, B_1_ or B_2_, with and without other variables, results in optimal performance for T. Thus there are 16 variable sets that can lead to optimal performance in the example. For an extensive discussion connecting marker/feature selection optimality, prediction and causal discovery see^26,27^.

#### Theoretical Analysis 1: Random feature selection and optimal predictive performance as a function of a number of Markov boundaries in data and their equivalence relations

As previously mentioned, if a selected feature set contains a MB(T), the best predictive model for T fitted with this feature set will achieve optimal performance (Supplementary Document 1, Remark 1 & 2). Therefore, the probability of a randomly selected feature set **S** resulting in optimal performance is the same as the probability of a randomly selected feature set of size |**S**| containing a MB(T) (i.e., it is a MBL(T)). The latter probability is affected by the total number of variables, the selected feature set size |**S**| and the equivalence structure of the MB(T)s (i.e., the composition of MBC-TIE sets, **Figure2**, **Supplementary Document 1** Definition 3&4). Three interdependent components contribute to the equivalence structure: the number of MBs, the size of each MB, and the variables involved in each MB. **Theorem 1** provides mathematical formulas for computing the probability of randomly selecting a MBL(T) of a specific size. To provide intuition, **Figure 3A and B** illustrates how different MB(T) equivalence structures affect the probability of randomly obtaining a MBL(T) of size 3. For a fixed equivalence structure, the probability of selecting a MBL(T) increases monotonically with variable set size |**S**|, and asymptotes at probability=1 (Figure 3B; **Theorem 2**). Trivially, the full set of variables is a MBL(T).

**Figure 3.**
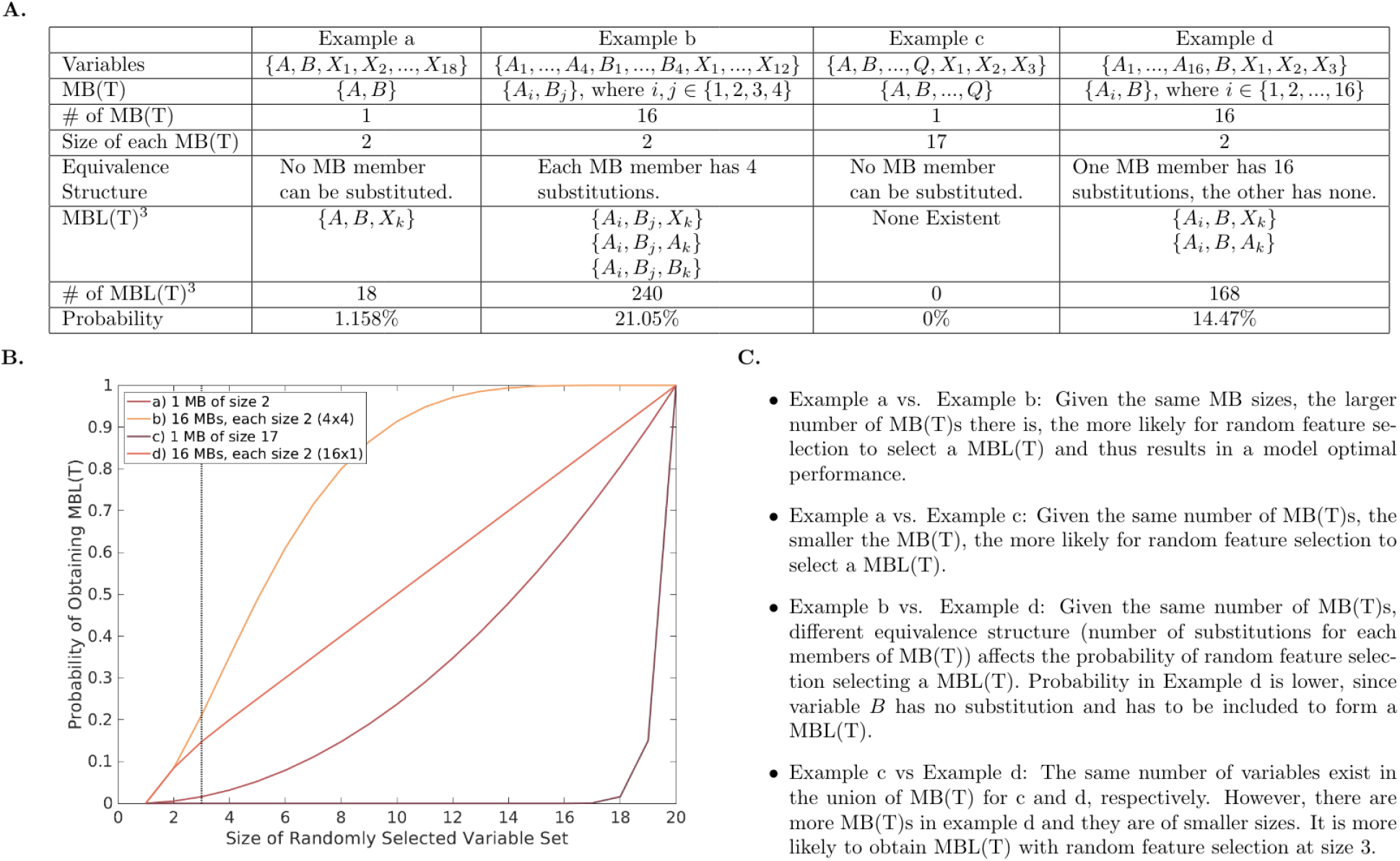
Relationship between Markov boundary equivalence structure and performance of random feature selection. Figure 3 shows four example distributions with different equivalence structures defined by three interdependent factors: the number of Markov Boundaries (MBs), the sizes of each MB, and the variables involved in each MB. They interact non-linearly and jointly influence the probability of random feature selection obtaining a Markov Blanket (MBL(T)) and thus resulting in optimal performance. Precise calculation of the probabilities is specified in **Theorem 1** (Supplementary Document 1). Panel A lists 4 examples of distributions with 20 variables for predicting their respective phenotype target of interest T with different equivalence structures. Examples (a) and (c) contain one unique MB(T) each, Examples (b) and (d) contain multiple MB(T)s. Due to the different equivalence structures, the number of Markov Blankets of size 3 (row 6 of table, denoted as MBL(T)^3^) differ across the 4 example distributions. As a result, the probability of randomly selecting a MBL(T) of size 3 differs drastically for the 4 distributions (row 8 of table). Panel B shows the probability of obtaining a MBL(T) via random feature selection for each example in A at when randomly selecting variable sets of different sizes. Panel C compares the differences in equivalence structure between different example distributions in A.

#### Theoretical Analysis 2: Structure and limitations of signature better-than-random tests of statistical significance

Fundamentally any better-than-random test of statistical significance must compare (by means of permutation or asymptotic strategies): (1) the predictive performance produced by a model built based on features selected using an algorithmic feature selection method or selected by experts, or a hybrid selection approach, with (2) the distribution of predictive performance produced by a model built based on randomly selecting features with the same feature set size (i.e., the null). In other words, predictivity is the test statistic (while controlling for parsimony). The p-value of the better-than-random test is computed as the proportion of random feature sets that perform better than that of the algorithmic (or human) feature selection. If one abides by common statistical practice, then statistical significance would occur when the p-value is ≤ α =0.05 (i.e., probability of false positives) or other suitable value for α. Because an asymptotic theory of distribution of predictive performance produced by a model built based on randomly selecting features with the same feature set size does not exist, we employ a permutation test as is common practice for testing properties of classifiers^28,29^.

**Theorem 3** establishes that the Markov Boundary in Faithful distributions has the smallest p-value among all sets with the same size and among all sets with optimal performance, and all variable sets of larger sizes. The p-value of MB is significant in most Faithful distributions. Notably, however statistical significance by the better-than-random test cannot be achieved in some distributions by any algorithm or experts, even when the feature set performs optimally (i.e. selects MB(T)) and even in the sample limit (**Theorem 4, Corollary 4.1**). E.g, in Figure 3 Example b, where multiple MB(T)s are present, 21.05% of the randomly selected feature sets of size 3 are MBL(T)s, leading to optimal performance, hence any feature selector choosing 3 features will not be statistically significantly different than random regardless of sample size.

#### Theoretical Analysis 3: Relationship between the causal significance of marker panels/signatures and their p-value for randomness

Can statistical significance from the better-than-random test be used to indicate causal significance (i.e., biological mechanistic relevance) of a feature set? **Theorem 5**, provides sufficient conditions under which the statistical significance of the better-than-random test guarantees causal significance. In Faithful distributions and without unmeasured confounders (aka “causal sufficiency”13), there is a direct correspondence between the variable set that produces optimal predictive performance, i.e. the unique MB(T), and the local causal neighborhood (“pathway”) of T, which consists of the direct causes, direct effects and direct causes of the direct effects. Therefore, in Faithful distributions with causal sufficiency, the local causal pathway of T, *and only that*, will result in statistical significance with the minimum p-value among all marker selectors returning sets of equal size or larger to the maximally parsimonious optimally predictive set. Unfortunately, **Theorem 5** also proves that the link between statistical significance and causality breaks down in TIE distributions, where only one out of the many MB(T)s corresponds to the local causal neighborhood of T. Finally in TIE distributions the local causal pathway, *even in the absence of unmeasured confounding*, is not guaranteed to be statistically significant in the better-than-random test.

### Revisiting Empirical Findings Using the Theoretical Framework

#### Empirical results closely track the theoretical analyses

Figure 1 (a) shows empirically that as the number of randomly selected features increases, the predictive performance of random feature selection increases. This is consistent with the monotonicity proof in **Theorem 2**. Further, as expected from **Theorems 1, 3, and 4**, given optimal or near-optimal predictive performance, the most parsimonious algorithm GLL-OPT more often outperforms random feature selection compared to T-fdr, because the latter algorithm systematically selects larger feature sets (**Table 1**). Further, when predictive performance is suboptimal, the better-than-random test can still be significant (**Table 1**, performance of RFVS).

#### Empirical “dissection” Of Markov boundary equivalence classes shows close correspondence between the theory and empirical testing of random feature selection

To test the correspondence of the formulas in **Theorem 1** and random feature selection in real-world data, we algorithmically extracted all MB(T)s in two gene expression datasets Alon^30^(|**V**|=2000) and Singh^31^(|**V**|=12600) using the TIE* algorithm^24^. The total number of MB(T)s identified were 9,501 and 6,058 respectively, providing empirical evidence for the presence of multiple MB(T)s in high-throughput molecular data (**Figure 4a, Supplementary table S8**). In addition, we show the close correspondence of probabilities calculated using **Theorem 1c** vs. empirically observed probability of obtaining MBs by random selection in these two datasets (**Figure 4c & 4d**). Prior literature hypothesized that in gene expression data from cancer tissues, algorithmic feature selection did not outperform random feature selection primarily due to the abundance of predictive genes related to cell proliferation^13^. To test this hypothesis, we mapped the variables in the identified MBs to known pathways^32^. Our results refute the proliferative markers hypothesis by showing genes from a variety of pathways (not just proliferative ones) can serve as optimal predictors (**Figure 4b**).

**Figure 4.**
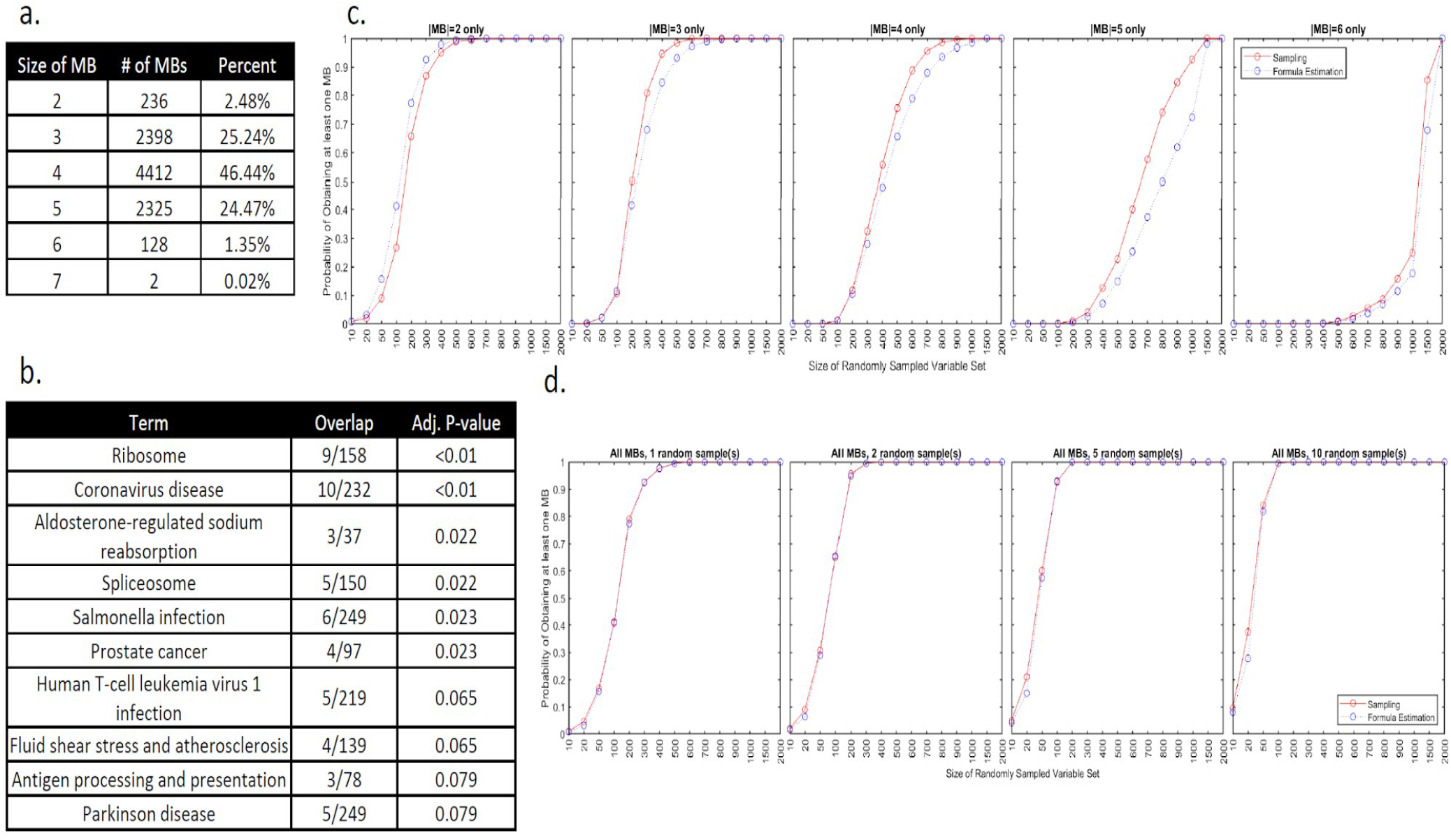
Dissection of Markov boundary equivalence classes in real-life datasets and testing the concordance between the expected and the observed performance of random feature selection. a) distribution of MB(T) sizes in the Alon dataset. b) The discovered MB(T) variables are involved in a multitude of gene pathways. c) The probability of randomly selecting a MB(T) of specific sizes estimated from our theoretical formula in Theorem 1c closely matches the one empirically estimated via random feature selection. d) The probability of randomly selecting a MB(T) of any size estimated from our theoretical formula in Theorem 1c closely matches the one empirically estimated via random feature selection.

#### Revisiting and comparison to Venet et al study findings

The Venet et al study^1^ findings which were focused on breast cancer bulk transcriptomics and a reference proliferation signature, at face value suggest that (1) random signatures correlate with the phenotype and (2) the proportion of published signatures were p-value matched or outperformed by random marker selection was 60%, hence much higher than the ones in our analyses. As shown in **Supplementary document 2**, while the main thesis of the Venet paper is correct, the specific estimates about random signature p-values were upward biased due to a combination of systematic and random biases in how the random testing was conducted, the data selection and the validation design.

## Discussion

The high predictivity of random signatures and markers, if not appropriately controlled and interpreted, threatens the rigor, (validity and completeness) of analyses of omics data and thus makes this issue particularly important across the translational spectrum from basic biology to translational and clinical research. The work presented here is therefore highly relevant to the work of numerous biomedical researchers from the biological, medical and computational disciplines.

Our approach combined extensive experimental analyses with deep theoretical (mathematical and computational) analyses. On the experimental testing front we conducted the largest empirical investigation of the subject (and one of the largest bioinformatics benchmark studies of any kind in the literature). The experiments involved fitting >120 million models (**Online Methods**) and established that the phenomenon of random marker and signature predictive optimality is more widespread than previously thought, in the sense of affecting practically all types of omics data, and mainstream discovery procedures.

Whereas experimental studies are valuable and insightful, they can cover only a small, finite set of cases. Theoretical analyses, on the other hand, can cover an infinite space of relevant possibilities. We therefore developed a comprehensive mathematical-computational framework that shows that the **root cause and sufficient and necessary condition for these “anomalous” signature behaviors** with respect to randomness is the existence, in each dataset, of a large equivalence class of optimally predictive feature sets *with respect to phenotypes of interest*. Various reasons lead to such equivalences.

The ***factors creating large Markov Boundary and Markov Blanket equivalence classes*** in omics data include: (1) *Engineered “redundancy” in the assays so that* the same *biological entity is measured in multiple parts of omics assays*^33^. (2) *“Superfluous signal”* because of s*ystem-level mismatches between response variables and predictor variables (i.e., the response variable, e.g., cancer vs normal phenotypes, encapsulates numerous more granular molecular processes each of which corresponds to different variables and is sufficient for the detection of macro states).* (3) *Variable measurement redundancies due to incomplete domain theory. (4) Small sample sizes/Limited statistical power*, Future work is warranted for deeper exploration of these and other factors that lead to large equivalence classes of optimal predictors. Note that factor #4 relates to insufficient power to show that minuscule predictivity differences are statistically significantly different (and thus the equivalences will vanish in the large sample). Importantly, however, the other factors #1-3 lead to distributions where the equivalences are structural and not due to small sample size, and thus *will not vanish in the large sample size*.

The theoretical framework encompasses precise definitions of factors involved, feasibility and impossibility theorems, and analytical formulas. The formulas compute the probability that a random signature achieving optimal phenotypic predictivity as a function of 1 user-controlled parameter (# of variables selected at random) and 2 (testable) data-dependent ones (structure of MB equivalence classes and the total number of variables in the dataset). The theorems establish under what conditions algorithmic marker selection will be statistically significantly different than random, under what conditions researchers can link the p-values of tests of randomness with predictive optimality, parsimony and mechanistic interpretation of signatures, and other properties.

Notably, our theoretical analyses cover *all possible algorithms regardless of what is their design*, because they reduce the infinite space of algorithms to a small number of identically-behaving algorithm types defined by the sufficient parameters of predictive accuracy and parsimony of their output. For the same reason, they apply equally well to human expert knowledge-derived signatures. *Thus, the results presented will not change if different signature discovery algorithms, human expert criteria, or background knowledge are introduced in the future*.

Importantly, our findings challenge the conventional “statistics 101” wisdom of conducting a test of statistical significance against the null hypothesis (i.e., signature not different than random signature selection) and ignoring signatures for which the null is not rejected. The theorems also show that in some omics distributions there *cannot possibly exist algorithms* or knowledge-driven procedures that will always (or in some cases ever) pass such a test, even in the sample limit.

A second round of empirical testing tested the mathematical predictions of the theory, and confirmed that there is a very close correspondence between the predictions of the theory and the actual empirical findings in both real data and in simulated data designed to test the theory.

Whereas the existence of large signature equivalence classes in omics data was demonstrated before^8,24,25^, here for the first time in the literature we present a **dissection of the equivalence class of biomarkers** in 2 datasets comprising (i) extraction of the full equivalence class’ member signatures and (ii) functional annotation via mapping them to biological pathways. This analysis shows that published explanations of random signature accuracy based on specific biological functional roles are incorrect^13^. We establish that random signature optimal predictive accuracy is not linked to any particular biological function (e.g., proliferation in cancer).

***MB equivalence classes can be (and are typically) invisible due to researchers’ lack of familiarity with them, and with equivalence class discovery algorithms***^24^. Without going into details, we also note that using collinearity as a criterion for target information equivalence is incorrect and will not work for discovery of MB and MBL equivalence classes.

### Implications and recommended good modeling and analysis practices

Good modeling practices in this particular area are essentially absent from the literature. Few studies^13,34^ addresses the issue but frames guidance based on assumptions that the present work proved to be inappropriate. We thus codify in **Table 2 proposed** good practices, informed by our experimental and theoretical findings. These GMPs address: prediction-focused studies, mechanism-focused studies and algorithm development efforts.

**Table 2.**
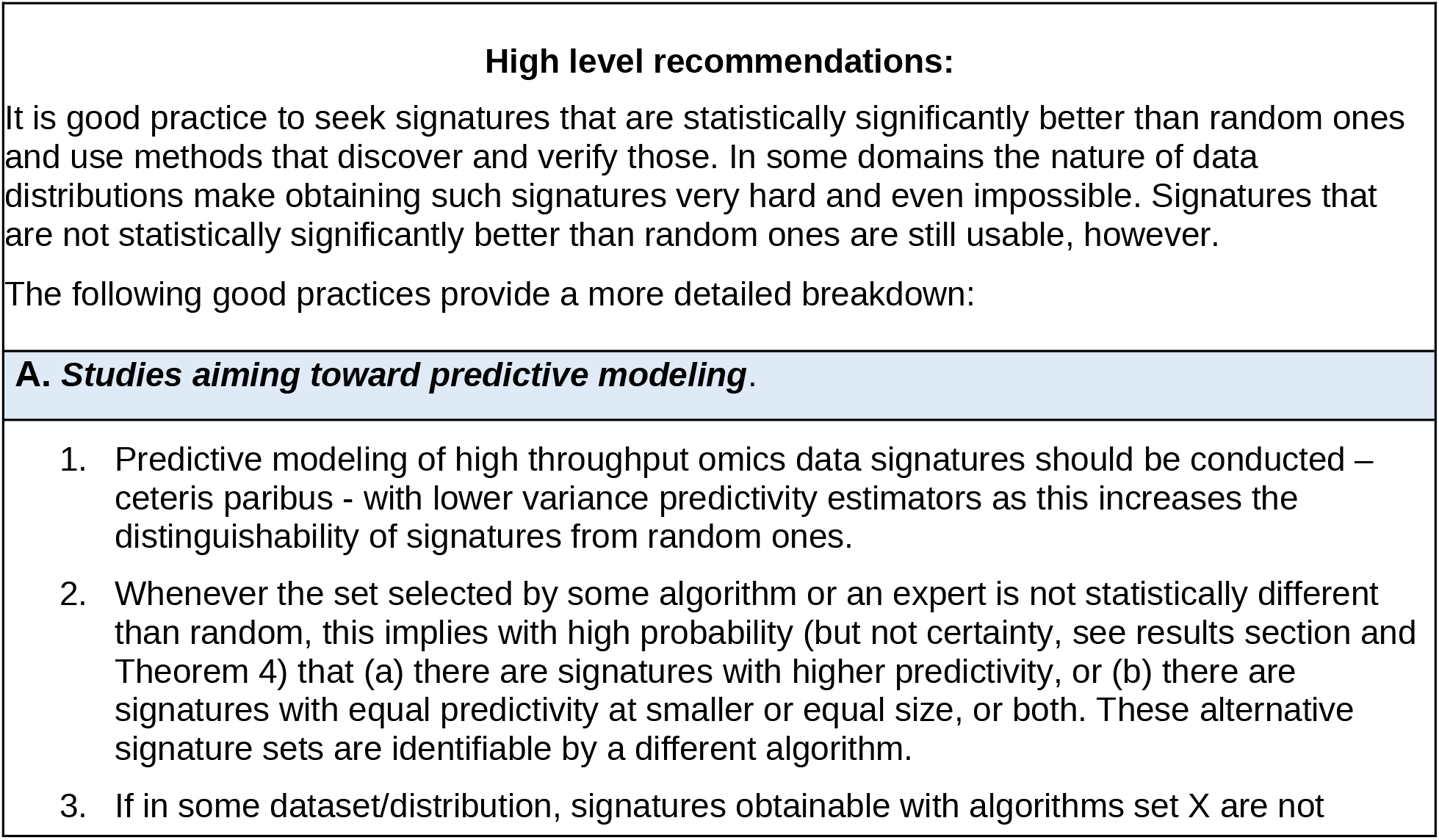
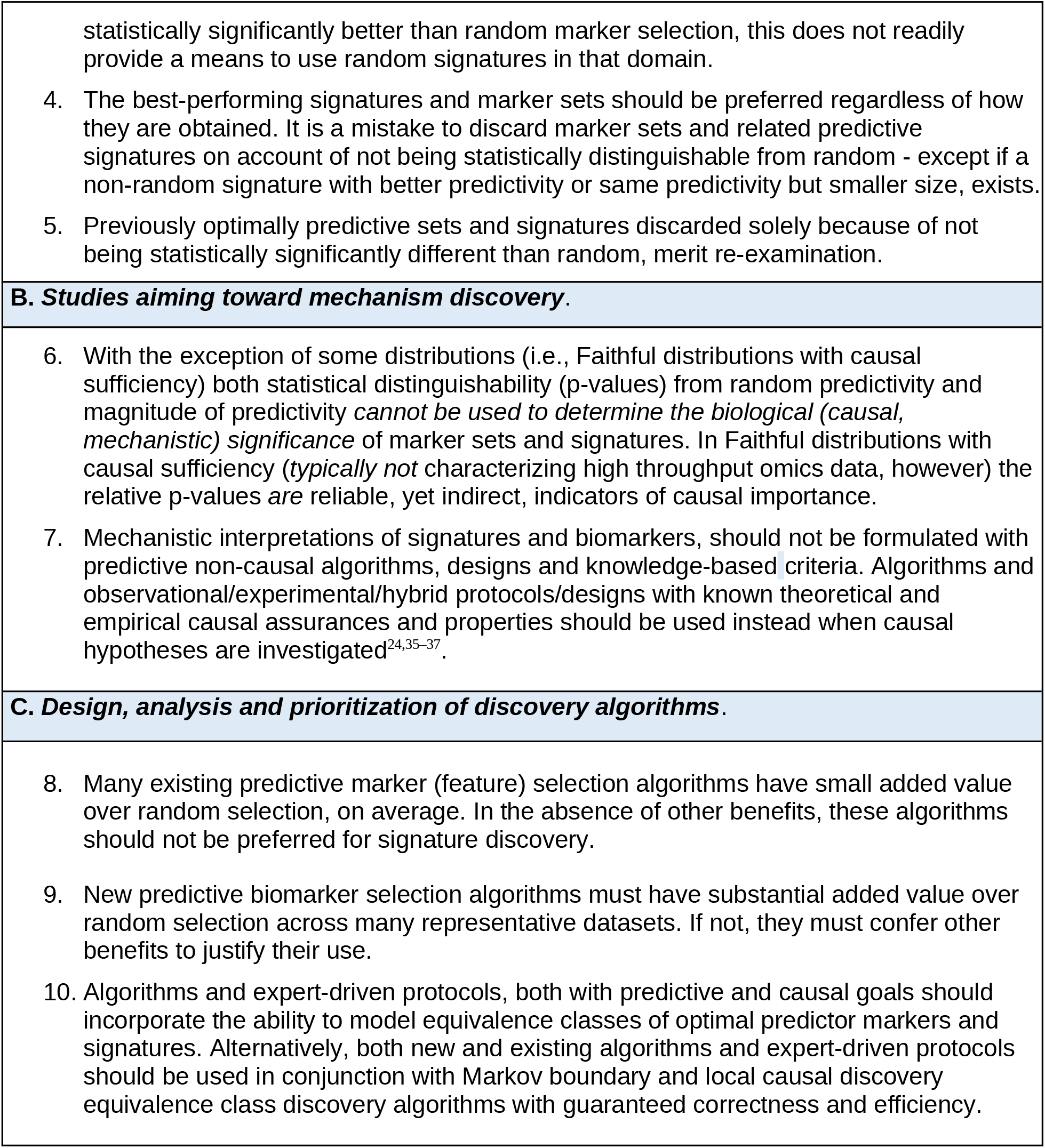
Good Practices for dealing with random predictive optimality in omics (and similar data) marker/signature study design and interpretation.

**In conclusion**, the existence of very large equivalence classes of optimally predictive feature sets, is a characteristic structural property of omics data, it does not vanish in large sample sizes, and it is exacerbated in small sample settings. This property creates “anomalous” behaviors of omics data with respect to randomness and these have implications for the design, analysis and interpretation of omics studies. We developed and tested a comprehensive methodological framework for studying and understanding the behavior of random molecular signatures and biomarkers and a corresponding good practices roadmap for dealing with randomness in omics data in order to avoid related common over- and under-interpretation pitfalls. These results apply to both algorithmic and human expert biomarker and signature discovery. Although omics data are replete with equivalent optimal predictors, the results of the present study extend to all types of distributions with similarly large equivalence classes of optimal predictors.

## Supporting information

SupplementaryDocument1_Proofs

Detailed_Methods

## Acknowledgement

We thank the SenNet Working group on Good Practices: Analytical Methods and Modeling (GP-AMM) and the SenNet Steering Committee for feedback on this work. The authors acknowledge the Minnesota Supercomputing Institute (MSI) at the University of Minnesota for providing resources that contributed to the research results reported within this paper.

## Footnotes

a We will use “biological marker” (or simply “marker”) to denote any molecular entity in omic data that is studied in the context of its predictive or causal relationship to phenotypes (e.g., biological function, disease or other) of interest. The term “biomarker” is within our scope but has a narrower interpretation since it typically relates to proxies of outcomes or indicators of side effects of therapeutic interventions^9,10^.

b In the context of the present report, a signature is a set of markers but also implies the corresponding mathematical or computational model(s) that take as input the biological markers’ values and output the most accurate predictions that the data allows with regards to the phenotype of interest. Such models can be developed with standard model fitting procedures using classifiers and modeling protocols with empirical and theoretical optimal performance^11,12^.

## Reference

1. Casella, G. et al. Transcriptome signature of cellular senescence. Nucleic Acids Res. 47, 7294–7305 (2019).

2. Gorgoulis, V. et al. Cellular Senescence: Defining a Path Forward. Cell 179, 813–827 (2019).

3. Wechter, N. et al. Single-cell transcriptomic analysis uncovers diverse and dynamic senescent cell populations. Aging 15, 2824–2851 (2023).

4. Li, S. et al. Advancing biological understanding of cellular senescence with computational multiomics. Nat. Genet. 57, 2381–2394 (2025).

5. Carlson, J. J. & Roth, J. A. The impact of the Oncotype Dx breast cancer assay in clinical practice: a systematic review and meta-analysis. Breast Cancer Res. Treat. 141, 13–22 (2013).

6. Milbury, C. A. et al. Clinical and analytical validation of FoundationOne®CDx, a comprehensive genomic profiling assay for solid tumors. PloS One 17, e0264138 (2022).

7. Cheng, D. T. et al. Memorial Sloan Kettering-Integrated Mutation Profiling of Actionable Cancer Targets (MSK-IMPACT): A Hybridization Capture-Based Next-Generation Sequencing Clinical Assay for Solid Tumor Molecular Oncology. J. Mol. Diagn. JMD 17, 251 –264 (2015).

8. Kraus, V. B. et al. Select Small Non-Coding RNAs Are Determinants of Survival in Older Adults. Aging Cell 25, e70403 (2026).

9. Kraus, V. B. et al. Proposed study designs for approval based on a surrogate endpoint and a post-marketing confirmatory study under FDA’s accelerated approval regulations for disease modifying osteoarthritis drugs. Osteoarthritis Cartilage 27, 571–579 (2019).

10. FDA-NIH Biomarker Working Group. BEST (Biomarkers, EndpointS, and Other Tools) Resource. (Food and Drug Administration (US), Silver Spring (MD), 2016).

11. Kearns, M. J., Schapire, R. E. & Sellie, L. M. Toward efficient agnostic learning. in Proceedings of the fifth annual workshop on Computational learning theory 341–352 (1992).

12. Statnikov, A., Aliferis, C. F., Tsamardinos, I., Hardin, D. & Levy, S. A comprehensive evaluation of multicategory classification methods for microarray gene expression cancer diagnosis. Bioinforma. Oxf. Engl. 21, 631–643 (2005).

13. Venet, D., Dumont, J. E. & Detours, V. Most Random Gene Expression Signatures Are Significantly Associated with Breast Cancer Outcome. PLOS Comput. Biol. 7, e1002240 (2011).

14. Daniel, W. W. & Cross, C. L. Biostatistics: A Foundation for Analysis in the Health Sciences. (John Wiley & Sons, 2018).

15. van ’t Veer, L. J., et al. Gene expression profiling predicts clinical outcome of breast cancer. Nature 415, 530–536 (2002).

16. Shaughnessy, J. D., Jr et al. A validated gene expression model of high-risk multiple myeloma is defined by deregulated expression of genes mapping to chromosome 1. Blood 109, 2276–2284 (2006).

17. Zou, H. & Hastie, T. Regularization and variable selection via the elastic net. J. R. Stat. Soc. Ser. B Stat. Methodol. 67, 301–320 (2005).

18. Saul, D. et al. A new gene set identifies senescent cells and predicts senescence-associated pathways across tissues. Nat. Commun. 13, 4827 (2022).

19. Heng, Y. J. et al. The molecular basis of breast cancer pathological phenotypes. J. Pathol. 241, 375–391 (2017).

20. Mohammed, H. et al. Progesterone receptor modulates ERα action in breast cancer. Nature 523, 313–317 (2015).

21. Braga-Neto, U. M. & Dougherty, E. R. Is cross-validation valid for small-sample microarray classification? Bioinformatics 20, 374–380 (2004).

22. Meek, C. Strong completeness and faithfulness in Bayesian networks. in Proceedings of the Eleventh conference on Uncertainty in artificial intelligence 411–418 (Morgan Kaufmann Publishers Inc., San Francisco, CA, USA, 1995).

23. Pearl, J. Causal inference in statistics: An overview. Stat. Surv. 3, 96–146 (2009).

24. Statnikov, A., Lytkin, N. I., Lemeire, J. & Aliferis, C. F. Algorithms for Discovery of Multiple Markov Boundaries. J. Mach. Learn. Res. JMLR 14, 499–566 (2013).

25. Statnikov, A. & Aliferis, C. F. Analysis and Computational Dissection of Molecular Signature Multiplicity. PLOS Comput. Biol. 6, e1000790 (2010).

26. Tsamardinos, I. & Aliferis, C. F. Towards Principled Feature Selection: Relevancy, Filters and Wrappers. in AISTATS (2003).

27. Simon, G. & Aliferis, C. An Appraisal and Operating Characteristics of Major ML Methods Applicable in Healthcare and Health Science. in Artificial Intelligence and Machine Learning in Health Care and Medical Sciences: Best Practices and Pitfalls (eds Simon, G. J. & Aliferis, C.) (Springer, Cham (CH), 2024).

28. Good, P. Permutation Tests: A Practical Guide to Resampling Methods for Testing Hypotheses. (Springer Science & Business Media, 2013).

29. Ojala, M. & Garriga, G. C. Permutation tests for studying classifier performance. J. Mach. Learn. Res. 11, (2010).

30. Alon, U. et al. Broad patterns of gene expression revealed by clustering analysis of tumor and normal colon tissues probed by oligonucleotide arrays. Proc. Natl. Acad. Sci. U. S. A. 96, 6745–6750 (1999).

31. Singh, D. et al. Gene expression correlates of clinical prostate cancer behavior. Cancer Cell 1, 203–209 (2002).

32. Chen, E. Y. et al. Enrichr: interactive and collaborative HTML5 gene list enrichment analysis tool. BMC Bioinformatics 14, 128 (2013).

33. Draghici, S., Khatri, P., Eklund, A. C. & Szallasi, Z. Reliability and reproducibility issues in DNA microarray measurements. Trends Genet. TIG 22, 101–109 (2006).

34. Beck, A. H. et al. Significance analysis of prognostic signatures. PLoS Comput. Biol. 9, e1002875 (2013).

35. Spirtes, P., Glymour, C. & Scheines, R. Causation, Prediction, and Search. (Springer-Verlag, New York, 1993).

36. Statnikov, A. et al. Ultra-scalable and efficient methods for hybrid observational and experimental local causal pathway discovery. J. Mach. Learn. Res. 16, 3219–3267 (2015).

37. Ma, S., Kemmeren, P., Aliferis, C. F. & Statnikov, A. An Evaluation of Active Learning Causal Discovery Methods for Reverse-Engineering Local Causal Pathways of Gene Regulation. Sci. Rep. 6, 22558 (2016).

