## SupplementaryDocument1_Proofs for "Optimal Predictivity of Random Signatures: Causes and Good Modeling Practices"

### Definitions and Proofs Related to Random Feature Selection

Jan 2026

#### Contents

**1 Introduction and Roadmap** **2**

**2 Notation** **2**

**3 Definitions and Relevant Prior Results** **2**

3.1 Markov Boundary, Markov Blanket, and Target Information Equivalence . . . . . . . . 2

3.2 Random Feature Selection . . . . . . . . . . . . . . . . . . . . . . . . . . . . . . . . . . . . . 4

**4 Theorems** **5**

#### List of Theorems

### 1 Introduction and Roadmap

This is a supplemental documents that includes related definitions and proofs for the theorems referenced in the main text of the paper, "Optimal predictivity and reproducibility of random omics biological markers and signatures: causes, consequences, and best practices".

Section 2 introduces notations. Section 3 contains definitions (some were defined in prior literature and some are introduced for the first time here) and prior results that are useful for background information for the new theoretical results presented in the current work. Section 3 are organized by definition of key concepts. For each definition, under the heading "Remark", we clarify the definition, introduce relevant previously proven theorems, and comment on their implications. Section 4 contains new theorems and their proofs.

### 2 Notation

Following convention, uppercase letters are used to represent variables (i.e. features), e.g. $A_1$ or $X$ or $T$. Lower case letters are used for parameters or indices, e.g. $m$ or $n$. Bold uppercase letters are used to represent set, excluding sets of sets, e.g. $\mathbf{S}$. We use uppercase letter in calligraphic font (\mathcal) to denote set of sets. We denote the set of all Markov boundaries for a target variable $T$ as $\mathcal{MB} = \{\mathbf{MB}_1, \mathbf{MB}_2, \ldots, \mathbf{MB}_i, \ldots\}$, where $\mathbf{MB}_i$ is a Markov boundary set. When indexing is not important, we use $\mathbf{MB}$ to refer to any Markov boundary set. When there is only one unique Markov boundary, i.e. $|\mathcal{MB}| = 1$, we refer to the unique Markov Boundary as $\mathbf{MB}$. Similarly, the set of all Markov blankets is denoted as $\mathcal{MBL} = \{\mathbf{MBL}_1, \mathbf{MBL}_2, \ldots \mathbf{MBL}_i, \ldots\}$, where $\mathbf{MBL}_i$ is a Markov blanket set. $\mathbf{MBL}$ is used to refer to any Markov blanket set when indexing is not important. The superscript for a set of sets is used to denote for sets of a specific size: $\mathcal{X}^n = \{\mathbf{X} \mid |\mathbf{X}| = n\}$. For example, $\mathcal{MBL}^3$ is a set of all Markov blanket sets of size 3.

### 3 Definitions and Relevant Prior Results

#### 3.1 Markov Boundary, Markov Blanket, and Target Information Equivalence

**Definition 1: Markov Boundary of $T$ [1].** *A Markov boundary of $T$, denoted as $\mathbf{MB}(T)$, is a minimal set of features with the following property: $\forall \mathbf{S} \subseteq \mathbf{V} \backslash \mathbf{MB}(T)$, $\mathbf{S} \perp T | \mathbf{MB}(T)$.*

**Remark 1: Commentaries and Prior Results Related to Markov Boundary.**

- *To note, the concept of Markov boundry is always in reference to a target variable $T$. For notational convenience, we use* $\mathbf{MB}$ *to denote* $\mathbf{MB}(T)$.

- $\perp$ *denotes independence, i.e.* $\forall \mathbf{S} \subseteq \mathbf{V} \backslash \mathbf{MB}$, $P(T, \mathbf{S}|\mathbf{MB}) = P(T|\mathbf{MB})P(\mathbf{S}|\mathbf{MB})$, *or equivalently,* $P(T|\mathbf{S}, \mathbf{MB}) = P(T|\mathbf{MB})$, *for any values of variables* $T$, $\mathbf{S}$, *and* $\mathbf{MB}$, *where* $P(\mathbf{MB}) > 0$. *This means,* $\mathbf{MB}$ *contains maximum information about* $T$ *among all subsets of* $\mathbf{V}$.

- *As noted in the definition,* $\mathbf{MB}$ *is minimal, i.e. removing any non-empty subset from* $\mathbf{MB}$ *will result in loss of information for* $T$.

- *Prior Results: Under the conditions that the learner that constructs the classification model can learn the distribution* $P(T|\mathbf{MB})$, *and that the loss function is such that perfect estimation of the probability distribution of* $T$ *is required. Models built with* $\mathbf{MB}$ *results in optimal performance for* $T$ *[2].*

- *Prior Results: In faithful distributions there is one unique* $\mathbf{MB}$ *for* $T$, *consisting of the parents, children, and parents of children of* $T$ *[3].*

**Definition 2: Markov Blanket [1].** *A variable set* $\mathbf{MBL}$ *where* $\mathbf{MB} \subseteq \mathbf{MBL}$ *is a Markov blanket.*

**Remark 2: Commentaries on Markov Blanket.**

- *Given the definition of* $\mathbf{MB}$ *and* $\mathbf{MBL}$, $\mathbf{MBL}$ *contains maximum information about* $T$ *over all subsets of* $\mathbf{V}$, *however it is not minimal.*

- *Similar to* $\mathbf{MB}$, *under the conditions that the learner that constructs the classification model can actually learn the distribution* $P(T|\mathbf{MBL})$, *and that the loss function is such that perfect estimation of the probability distribution of* $T$ *is required. Models built with* $\mathbf{MBL}$ *results in optimal performance for* $T$.

**Definition 3: Markov Boundary Conditional Target Information Equivalence (MBC-TIE).** *Feature sets* $\mathbf{S}_1$ *and* $\mathbf{S}_2$ *are Markov boundary conditional target information equivalent when there exist two Markov boundaries of* $T$, $\mathbf{MB}_1$ *and* $\mathbf{MB}_2$, *such that:* $\mathbf{S}_1 \subset \mathbf{MB}_1$, $\mathbf{S}_2 \subset \mathbf{MB}_2$, $\mathbf{MB}_1 = \mathbf{S}_1 + \mathbf{MB}_1 \cap \mathbf{MB}_2$, $\mathbf{MB}_2 = \mathbf{S}_2 + \mathbf{MB}_1 \cap \mathbf{MB}_2$. *These equations are referred to as substitution functions, since they specify that one can obtain*$\mathbf{MB}_2$ *by substituting* $\mathbf{S}_1$ *with* $\mathbf{S}_2$ *in* $\mathbf{MB}_1$ *and vice versa.*

**Remark 3: Commentaries on Multiple Markov Boundaries in a MBC-TIE.**

- *Prior Results: Under the conditions that the learner that constructs the classification model can actually learn the distribution* $P(T|\mathbf{MB}_i)$, *and that the loss function is such that perfect estimation of the probability distribution of* $T$ *is required. Models built with any* $\mathbf{MB}_i$, *results in optimal performance for* $T$ *[4].*

- *Given the definition of* $\mathbf{MB}$, $\forall \mathbf{MB}_i \in \mathcal{MB}$ *is minimal, i.e. removing any non-empty subset from* $\mathbf{MB}_i$ *will result in loss of information for* $T$.

- *The substitution functions also suggest that* $\mathbf{S}_1$ *is information equivalent with* $\mathbf{S}_2$ *with respect to* $T$ *given* $\mathbf{MB}_1 \cap \mathbf{MB}_2$. *Therefore, the substitution functions characterize the equivalence structure among the members of* $\mathcal{MB}$.

- *This definition is introduced by the authors for the first time. It is used in Theorem 1.*

**Definition 4: TIE Groups for One-to-one MBC-TIE.** *A one-to-one MBC-TIE is a special case of MBC-TIE that satisfies the following criterion: there exists a partition* $\Pi = \mathcal{TG} = \{\mathbf{TG}_1, \ldots, \mathbf{TG}_g\}$ *of the union of all Markov boundaries, such that their unordered Cartesian product results in the set of all Markov boundary sets, i.e.* $\mathcal{MB} = \bigotimes_{i=1}^{k} \mathbf{TG}_i$. *Each cell of the partition, i.e.* $\mathbf{TG}_i$, *is a one-to-one MBC-TIE group.*

**Remark 4: Commentaries One-to-one MBC-TIE group.**

- *$\mathcal{TG}$ denotes the set of all MBC-TIE groups. It is a partition of the union set of all Markov boundaries, which means any variable that is in the union set of all Markov boundaries appears in exactly one $\mathbf{TG}_i \in \mathcal{TG}$.*

- *Each Markov boundary contains one and only one member from each $\mathbf{TG}_i$. The set of all Markov boundaries $\mathcal{MB}$ can be constructed from the MBC-TIE groups.$|\mathcal{MB}| = \prod_{i=1}^{g} \mathbf{N}_{TG}$ and $|\mathbf{MB}_i| = g$.*

- *$\mathbf{N}_{\mathcal{TG}} := \{|\mathbf{TG}_1|, \ldots, |\mathbf{TG}_g|\}$, is the set that contains the number of variables in each MBC-TIE group.*

- *$\forall i$, $\forall X, Y \in \mathbf{TG}_i$, $X \perp T|\{Y, \cup\mathcal{MB}\backslash\mathbf{TG}_i\}$, and $Y \perp T|\{X, \cup\mathcal{MB}\backslash\mathbf{TG}_i\}$. This means any members of a one-to-one MBC-TIE group contains the same unique information regarding $T$ conditioning on the union of all Markov boundaries excluding the variables from the same MBC-TIE group.*

- *This definition is introduced by the authors for the first time. It is used in Theorem 1.*

**Example 1.** *We show in this example the correspondence between Markov boundaries with one-to-one MBC-TIE and their MBC-TIE groups. For the following $\mathcal{MB}$,*

$$\mathcal{MB} = \{\{A_1, B_1\}, \{A_1, B_2\}, \{A_1, B_3\}, \{A_2, B_1\}, \{A_2, B_2\}, \{A_2, B_3\}\}. \tag{1}$$

*The following partition of $\cup_i \mathbf{MB}_i$ exactly reproduce $\mathcal{MB}$ via their unordered Cartesian product:*

$$\mathbf{TG}_1 = \{A_1, A_2\} \tag{2}$$

$$\mathbf{TG}_2 = \{B_1, B_2, B_3\}, \tag{3}$$

*In contrast, for a different $\mathcal{MB}$,*

$$\mathcal{MB} = \{\{A_1, B_1\}, \{A_1, B_2\}, \{A_1, B_3\}, \{A_2, B_1\}, \{A_2, B_2\}\}. \tag{4}$$

*No partition of $\cup_i \mathbf{MB}_i$ exactly reproduce $\mathbf{MB}$ with unordered Cartesian product. So this $\mathcal{MB}$ does not have a one-to-one MBC-TIE structure.*

##### 3.2 Random Feature Selection

**Definition 5: Metric for predictive performance [2].** *$M : \langle \mathbf{C}, \mathbf{S}, T, D\rangle \to \mathbb{R}$ is a performance metric for predicting a target variable of interest $T$, where $D$ is a sample of observations over the variable set $\mathbf{V}$. $D \downarrow \{\mathbf{S}, T\}$ denotes the projection of $D$ to $\{\mathbf{S}, T\}$, where $\mathbf{S} \subseteq \mathbf{V}$. $\mathbf{C}$ is a set of classification learners. A classification learner $C$ takes $D \downarrow \{\mathbf{S}, T\}$ and outputs a prediction model for $T$. We assume $\forall \mathbf{S} \subseteq \mathbf{V}$, at least one member of the set of learners $\mathbf{C}$ can capture all the information contained in $\mathbf{S}$ regarding $T$.*

**Remark 5: Note on Performance Metric.**

- *To simplify notation, we define $M(\mathbf{S}, T, D) := \text{maximize}_{\mathbf{C}}(M(\mathbf{S}, \mathbf{C}(T, D \downarrow \mathbf{S})))$, denoting the value of metric $M$ maximized over the set of classifier learners $\mathbf{C}$ for predicting $T$ using feature set $\mathbf{S} \subseteq \mathbf{V}$ and data sample $D$.*

**Definition 6: Random Feature Selection.** *Random Feature Selector $R^k$ selects a variable set $\mathbf{S}$ of size $k$ , where $\forall V_i \in \mathbf{S}$ is selected without replacement from $\mathbf{V}$ with probability $\frac{1}{|\mathbf{V}|}$. The feature sets selected by the random feature selection are termed random feature sets.*

**Remark 6: Commentary on Random Feature Selection.**

- *The performance of the random feature selection is a uniform random sample from* $P_k(M(\mathbf{S}^k, T, D))$*, where* $P_k(M(\mathbf{S}^k, T, D))$ *denote the probability mass function of* $M(\mathbf{S}^k, T, D)$*, where* $\mathbf{S}^k := \{\mathbf{S} \subseteq \mathbf{V} \mid |\mathbf{S}| = k\}$*, i.e.,* $\mathbf{S}^k$ *denotes the set of subsets of* $\mathbf{V}$ *with size* $k$*.*

**Definition 7: A Better-than-random Test.** *To test the statistical significance for a feature set* $\mathbf{S}_{\mathrm{F}}$ *against randomly selected feature sets of the same size, reject* $H_0 : M(\mathbf{S}_{\mathrm{F}}, T, D) \leq M(\mathbf{S}_{R^{|\mathbf{S}_{\mathrm{F}}|}}, T, D)$ *when,*

$$p := 1 - \sum_{M < M(\mathbf{S}_{\mathrm{F}}, T, D)} P_{|\mathbf{S}_{\mathrm{F}}|}(M(\mathbf{S}^{|\mathbf{S}_{\mathrm{F}}|}, T, D)) < \alpha, \tag{5}$$

*where* $\mathbf{S}_{\mathrm{F}}$ *denotes a feature set selected by feature selection method* F. $\mathbf{S}_{R^{|\mathbf{S}_{\mathrm{F}}|}}$ *denotes a feature set selected by the random feature selector selecting the same number of variables as* $\mathbf{S}_{\mathrm{F}}$*.* $\alpha$ *is a desired maximum critical value for type I error.*

**Remark 7: Commentaries on the Better-than-random Test.**

- $P_{|\mathbf{S}_{\mathrm{F}}|}(M(\mathbf{S}^{|\mathbf{S}_{\mathrm{F}}|}, T, D))$ *is the null distribution of the better than random test.*
- $\sum_{M < M(\mathbf{S}_{\mathrm{F}}, T, D)} P_{|\mathbf{S}_{\mathrm{F}}|}(M(\mathbf{S}^{|\mathbf{S}_{\mathrm{F}}|}, T, D))$ *is the cumulative distribution of the null distribution.*
- *The better-than-random test is statistically significant when the percentage of random feature sets that have greater or the same performance compared to* $\mathbf{S}_{\mathrm{F}}$ *is smaller than* $\alpha$*.*
- *Commentary: Because the form of the sampling distribution of* $M(\mathbf{S}, T, D)$ *over all feature selectors and classifiers is a priori unconstrained, we test the hypothesis with an exact test.*
- *This form of better-than-random test has been described in the applied literature before. We provide a formal definition so its properties can be analyzed.*

### 4 Theorems

**Theorem 1: Probability of obtaining at least one Markov Boundary with random feature selection of size** $n$**.**

a) *In all distributions over a variable set* $\mathbf{V}$*:*

$$P(n; |\mathbf{V}|) = \frac{1}{\binom{|\mathbf{V}|}{n}} |\mathcal{MBL}^n|, \tag{6}$$

*where* $\mathcal{MBL}^n$ *is the set of all Markov blankets of size* $n$*.*

b) *In the most general case of unconstrained Markov boundary conditional target information equivalence structure (MBC-TIE, see Definition 3) among members of Markov boundary equivalence class:*

$$P(n; |\mathbf{V}|, \mathcal{MB}) = \frac{1}{\binom{|\mathbf{V}|}{n}} \sum_{\varnothing \neq \mathcal{S} \subseteq \mathcal{MB}} (-1)^{|\mathcal{S}|+1} \binom{|\mathbf{V}| - |\cup \mathcal{S}|}{n - |\cup \mathcal{S}|}, \tag{7}$$

*where* $\mathcal{MB} = \{\mathbf{MB}_1, \mathbf{MB}_2, \ldots, \mathbf{MB}_m\}$ *is the set of all Markov boundaries.*[a]

[a]We use the common definition of $\binom{n}{k} = 0$ when $k < 0$ or $k > n$.

c) *In a more special case of one-to-one MBC-TIE distributions (see Definition 4)*

$$P(n; |\mathbf{V}|, \mathbf{N}_{\mathcal{TG}}) = \sum_{\mathbf{J} \subseteq \{1,\ldots,k\}} (-1)^{|\mathbf{J}|} \binom{|\mathbf{V}| - \sum_{j \in \mathbf{J}} |\mathbf{TG}_j|}{n} \tag{8}$$

*where* $\mathbf{N}_{\mathcal{TG}} = \{|\mathbf{TG}_1|, \ldots, |\mathbf{TG}_g|\}$*, i.e.* $\mathbf{N}_{\mathcal{TG}}$ *is a set of size* $g$*, each element represents the number of variables in a MBC-TIE group.*

*Proof Sketch*

Theorem 1 supplies formulas for computing the probability of obtaining at least one Markov boundary when randomly selecting $n$ features with the random feature selector $R^n$ (Definition 6). Under the conditions specified in Remark 1, models based on Markov boundary variables result in optimal performance. Therefore, this probability is also the probability that $R^n$ resulting in optimal performance. Part a) formula states that the probability of selecting at least one Markov boundary when random selecting $n$ variables is equal to the number of Markov blankets of size $n$ over the total number of variable sets of size $n$. This directly follows the definition of Markov blanket (Definition 2).

Counting the number of Markov blankets of size $n$ is simple when there is only one Markov boundary. Consider a Markov boundary of size $l$ ($l \leq n$). By padding $n-l$ variables to the Markov boundary, we can obtain Markov blankets of size $n$. There are $\binom{|\mathbf{V}|-l}{n-l}$ ways to obtain a Markov blanket of size $n$ from a Markov boundary of size $l$. However, when we have more than one Markov boundaries, we can not simply pad each Markov boundary to Markov blankets of size $n$ and sum up the number of Markov blankets obtained. This is because two different Markov boundaries can be padded to the same Markov blanket [b].

Part b) and c) computes the number of Markov blankets of size $n$ under the unconstrained MBC-TIE structure (high computation complexity) and the one-to-one MBC-TIE (much lower computation complexity) respectively leveraging the inclusion-exclusion principle [5] to handle duplicate counting.

*Proof of Theorem 1a.* The number of possible ways to choose $n$ variables from a total of $|\mathbf{V}|$, is the binomial coefficient $\binom{|\mathbf{V}|}{n}$. The set of variables of size $n$ that includes at least one Markov boundary—hence has optimal performance—are the set of Markov blankets with size $n$, denoted by $\mathcal{MBL}^n$. So $P = |\mathcal{MBL}^n| / \binom{|\mathbf{V}|}{n}$. □

*Proof of Theorem 1b.* We compute $|\mathbf{MBL}^n|$ in the general case where the structure of MBC-TIE is unconstrained. Markov blanket of size $n$ is obtained by padding $n-l$ variables to a Markov boundary of size $l$, ($l \leq n$). There are $\binom{|\mathbf{V}|-l}{n-l}$ ways to pad a single Markov boundary of size $l$ into Markov blankets of size $n$.// When multiple Markov boundaries are present, we can pad each of the Markov boundaries to obtain a Markov blanket of size $n$. We denote the set of all sets of size $n$ that contains $\mathbf{MB}_i$ with $\mathcal{A}_i^n$. The number of ways to pad Markov boundary $\mathbf{MB}_i$ into Markov blankets of size $n$ is,

$$|\mathcal{A}_i^n| = \binom{|\mathbf{V}| - |\mathbf{MB}_i|}{n - |\mathbf{MB}_i|}. \tag{9}$$

Summing $|\mathcal{A}_i^n|$ over each $\mathbf{MB}_i$ would result in counting some Markov blankets more than once. Since sometimes two different Markov boundaries can be padded into the same Markov blankets. To correct for the over counting, we apply the inclusion-exclusion principle of combinatorics,

$$\left| \bigcup_{i=1}^{M} \mathcal{A}_i^n \right| = \sum_{\varnothing \neq \mathbf{J} \subseteq \{1,\ldots,M\}} (-1)^{|\mathbf{J}|+1} \left| \bigcap_{j \in \mathbf{J}} \mathcal{A}_j^n \right| \tag{10}$$

[b] Consider $\mathbf{V} = \{A, B, C, D\}$, $\mathcal{MB} = \{\{A, B\}, \{A, C\}\}$. Padding $\{A, B\}$ to size 3 results in $\{A, B, C\}$, and $\{A, B, D\}$. Padding $\{A, C\}$ to size 3 results in $\{A, B, C\}$ and $\{A, C, D\}$. $\{A, B, C\}$ will be counted twice if we simply sum the number of resulting Markov blankets over all the Markov boundaries. Consider it from another point of view, $\{A, B, C\}$ contains two Markov boundaries $\{A, B\}$ and $\{A, C\}$.

The left hand side of Equation (10) is the number of Markov blanket sets of size $n$, i.e. $\mathcal{MBL}^n$. Since the union operation removes the duplicate Markov blanket sets. The right hand side of Equation (10) is the inclusion-exclusion principle. Where we need to count the size of all intersection sets of $\mathcal{A}_j^n$. For example, the intersection set of $\mathcal{A}_i^n$ and $\mathcal{A}_j^n$ represents the Markov blanket of size $n$ that contains both $\mathbf{MBL}_i$ and $\mathbf{MBL}_j$, they are subtracted from Equation (10) to correct overcounting.

Now we expand the right hand side of Equation (10), levering the fact that the number of Markov blankets of size n that contain multiple Markov boundaries, e.g. $\mathcal{S} = \{\mathbf{MB}_i, \mathbf{MB}_j, \mathbf{MB}_k, \dots\}$, can be computed by $\binom{|\mathbf{V}|-|\cup\mathcal{S}|}{n-|\cup\mathcal{S}|}$, i.e., padding on top of variables that belongs to any MBs.

$$|\mathcal{MBL}^n| = \sum_{\varnothing \neq \mathcal{S} \subseteq \mathcal{MB}} (-1)^{|\mathcal{S}|+1} \binom{|\mathbf{V}| - |\cup \mathcal{S}|}{n - |\cup \mathcal{S}|} \tag{11}$$

Where the sum iterates over all non-empty subsets of $\mathcal{MB}$, the set of all Markov boundary sets. Plugging Equation (11) into Equation (6), we arrive at equation (7), completing the proof for Theorem 1b. □

**Example 2.** *We illustrate the above counting process with an example:*

$$\mathbf{V} = \{A_1, A_2, B_1, B_2, B_3, X_1, X_2, X_3\}. \tag{12}$$

$$\mathcal{MB} = \{\{A_1, B_1\}, \{A_1, B_2\}, \{A_1, B_3\}, \{A_2, B_1\}, \{A_2, B_2\}, \{A_2, B_3\}\}. \tag{13}$$

*We compute the number of Markov blankets of size $n = 3$ according to Theorem 1b.*
*For $|\mathcal{S}| = 1$ part of the sum we have,*

$$\sum_{\mathcal{S} \in \mathcal{MB}, |\mathcal{S}|=1} \binom{|\mathbf{V}| - |\cup \mathcal{S}|}{n - |\cup \mathcal{S}|} = \sum_{\mathbf{MB}_i \in \mathcal{MB}} \binom{|\mathbf{V}| - |\mathbf{MB}_i|}{n - |\mathbf{MB}_i|} = \binom{8-2}{3-2} \times 6 = 36, \tag{14}$$

*which is the sum over the number of ways to pad every Markov boundary into Markov blankets of size 3. But in Equation (14) we have counted some Markov blankets more than once. For example, the Markov blanket $\{A_1, B_1, A_2\}$ were counted both when we pad Markov boundary $\{A_1, B_1\}$ and $\{A_2, B_1\}$, since $\{A_1, B_1, A_2\}$ contains two Markov boundaries. To remove the duplicate counting, we need to subtract the number of Markov blankets of size 3 that contain 2 Markov boundaries: i.e. $|\mathcal{S}| = 2$,*

$$\begin{aligned} \sum_{\mathcal{S} \subset \mathcal{MB}, |\mathcal{S}|=2} \binom{|\mathbf{V}| - |\cup \mathcal{S}|}{n - |\cup \mathcal{S}|} &= \sum_{\mathbf{S}=\{\mathbf{MB}_i, \mathbf{MB}_j\}, i<j} \binom{|\mathbf{V}| - |\mathbf{MB}_i \cup \mathbf{MB}_j|}{n - |\mathbf{MB}_i \cup \mathbf{MB}_j|} \\ &= \binom{8-3}{3-3} \times 9 + \binom{8-4}{3-4} \times 6 = 9. \end{aligned} \tag{15}$$

*For this example, it is not possible to encounter a Markov blanket of size $n = 3$ that contains 3 or more Markov boundaries. Because in this example for any $|\mathcal{S}| \geq 3$ we have $|\cup \mathcal{S}| \geq 4$, which means $n - |\cup \mathcal{S}| < 0$ in any binomial coefficients in the sum.*

*Therefore, $|\mathcal{MBL}^3| = 36 - 9 = 27$ and $P = 27/\binom{8}{3} = 27/56 \approx 48.21\%$. So if we select $n = 3$ variables from $\mathbf{V}$ given by Equation (12), there is approximately 48% chance to get a feature set that contains at least one Markov boundary set specified in Equation (13).*

**Remark 8: Computational Cost of Equation (7).** *Note that given Equation (7), to compute the number of Markov blankets of size n, knowing the number of Markov boundaries and their sizes is not sufficient. We also need to be able to compute the size of the union of the Markov boundaries. Equation (7) applies to data with arbitrary MBC-TIE structure, which makes it general. However, this comes at a high computational cost. When the number of randomly selected features $n > |\cup_j \mathbf{MB}_j|$, the sum in Equation (7) runs over $2^{|\mathcal{MB}|} - 1$ subsets and requires $2^{|\mathcal{MB}|} - 1$ specification of sizes for union sets.*

*Proof of Theorem 1c.* In distributions that are one-to-one MBC-TIE (Definition 3), the expensive set union operation in equation (7) can be avoided and thus the computation of $P$ is greatly simplified.

Given MBC-TIE, it is easier to compute $|\mathcal{MBL}^n|$ by computing its complement, i.e. the number of sets of size $n$ that are not Markov blankets. This is because we need at least one member from each MBC-TIE group to form a Markov blanket[c]. In other words, any set of size $n$ that is not a Markov blanket does not contain any variables from at least one MBC-TIE.

We denote the set of all subsets of $\mathbf{V}$ (i.e. the power set of $\mathbf{V}$) as $\mathcal{P} = \{\mathbf{P}|\mathbf{P} \subseteq \mathbf{V}\}$. The subsets of $\mathbf{V}$ with size $n$ is $\mathcal{P}^n$. We denote the set of all subsets of size n that does not contain any variables from $\mathbf{TG}_i$ with $\mathcal{B}^n_{\hat{i}}$. We have,

$$\mathcal{MBL}^n = \mathcal{P}^n - \bigcup_{i=1}^{k} \mathcal{B}^n_{\hat{i}}, \tag{16}$$

$\bigcup_{i=1}^{k} \mathcal{B}^n_{\hat{i}}$ contains all the sets that are at least missing one MBC-TIE group. In other words, anything outside this union will have to have at least one member from each MBC-TIE group. To note: $|\bigcup_{i=1}^{g} \mathcal{B}^n_{\hat{i}}|$, the number of non-Markov blanket sets of size $n$ can not be computed by simply adding together $|\mathcal{B}^n_{\hat{i}}|$ over different $\mathbf{TG}_i$. This is because a specific non-Markov blanket set of size $n$ can miss variables from multiple different MBC-TIE groups. Therefore, $|\bigcup_{i=1}^{g} \mathcal{B}^n_{\hat{i}}|$ need to be computed leveraging the inclusion-exclusion principle. Considering the non-trivial case of $n \geq g$, to compute $|\mathcal{MBL}^n|$, we have:

$$|\mathcal{MBL}^n| = \binom{|\mathbf{V}|}{n} - \left|\bigcup_{i=1}^{g} \mathcal{B}^n_{\hat{i}}\right| \tag{17}$$

$$= \binom{|\mathbf{V}|}{n} - \sum_{\varnothing \neq \mathbf{J} \subseteq \{1,\ldots,k\}} (-1)^{|\mathbf{J}|+1} \left|\bigcap_{j\in\mathbf{J}} \mathcal{B}^n_{\hat{j}}\right| \tag{18}$$

$$= \binom{|\mathbf{V}|}{n} - \sum_{\varnothing \neq \mathbf{J} \subseteq \{1,\ldots,k\}} (-1)^{|\mathbf{J}|+1} \binom{|\mathbf{V}| - \sum_{j\in\mathbf{J}} |\mathbf{TG}_j|}{n} \tag{19}$$

$$= \sum_{\mathbf{J} \subseteq \{1,\ldots,k\}} (-1)^{|\mathbf{J}|} \binom{|\mathbf{V}| - \sum_{j\in\mathbf{J}} |\mathbf{TG}_j|}{n} \tag{20}$$

Equation (18) computes $|\bigcup_{i=1}^{g} \mathcal{B}^n_{\hat{i}}|$ using the inclusion-exclusion principle. Equation (19) applies the definition of $\mathcal{B}^n_{\hat{i}}$. Equation (20) simplifies Equation (19) by absorbing the $\binom{|\mathbf{V}|}{n}$ term into the sum (corresponding to adding back $\varnothing$) and thus leading to modification of the exponent of the sign term.

Lastly, using Theorem 1a, it follows that the probability of selecting at least one Markov boundary when randomly choosing $n$ variables out of $|\mathbf{V}|$ is as stated in equation (8). This completes the proof of Theorem 1c. □

**Example 3.** *To illustrate equation* (8) *with a concrete example, let,*

$$\mathbf{V} = \{A_1, A_2, B_1, B_2, B_3, X_1, X_2, X_3\}, \tag{21}$$

$$\mathbf{TG}_1 = \{A_1, A_2\}, \tag{22}$$

$$\mathbf{TG}_2 = \{B_1, B_2, B_3\}, \tag{23}$$

$$\mathbf{N}_{\mathcal{TG}} = \{2, 3\}. \tag{24}$$

*And given the MCB-TIE assumption of Theorem 1c,*

$$\mathbf{MB} = \mathbf{TG}_1 \otimes \mathbf{TG}_2 \tag{25}$$

$$= \{\{A_1, B_1\}, \{A_1, B_2\}, \{A_1, B_3\}, \{A_2, B_1\}, \{A_2, B_2\}, \{A_2, B_3\}\}. \tag{26}$$

[c]Note: $|\mathcal{MBL}^n| = 0$ for $n < k$

*Suppose we are selecting $n = 3$ variables out of $\mathbf{V}$. For the three selected variables to not contain a Markov boundary, the selected variables should not contain any variable in $\mathbf{TG}_1$, or $\mathbf{TG}_2$. The number of sets where selected variables do not contain variables in $\mathbf{TG}_1$, or $\mathbf{TG}_2$ are:*

$$\sum_{\mathbf{S}\subseteq\mathbf{N}_{\mathcal{TG}},|\mathbf{S}|=1} \binom{|\mathbf{V}| - \sum_{S_j\in\mathbf{S}} S_j}{n} = \binom{8-2}{3} + \binom{8-3}{3} = 30. \tag{27}$$

*Note that sets are neither in $\mathbf{TG}_1$, nor $\mathbf{TG}_2$ were counted twice. The number of these sets are computed when $|\mathbf{S}| = 2$,*

$$\sum_{\mathbf{S}\subseteq\mathbf{N}_{\mathcal{TG}}} \binom{|\mathbf{V}| - \sum_{S_j\in\mathbf{S}} S_j}{n} = \binom{8-(2+3)}{3} = 1. \tag{28}$$

*Subtracting equation (28) from equation (27) gives us the number of variable sets of size $n = 3$ that does not contain any Markov boundary. This subtraction corresponds to the alternating sum in equation (8),*

$$\frac{\binom{8}{3} + (-1)^1 \times 30 + (-1)^2 \times 1}{\binom{8}{3}} = \frac{27}{56} \approx 48.21\%. \tag{29}$$

**Remark 9: Computational Cost of Equation (8).** *The two sums in Equation (8) runs through $g2^{g-1}$ terms, where $g$ is the size of individual Markov boundaries. Under one to one MBC-TIE and for non-trivial cases, Equation (8) is able to compute the probability of obtaining Markov Blanket while random feature selection more efficiently compared to Equation (7), which has $2^{|\mathcal{MB}|} - 1$ terms to sum, and requires set union operations.*

**Remark 10: Relation to the statistical significance of the better-than-random test.** *The p-value of the better-than-random-test (Definition 7) for a feature set $\mathbf{S}_\mathrm{F}$ is the percentage of times that randomly selected variable sets of the same size performs better than $\mathbf{S}_\mathrm{F}$. When the probability of a randomly selected feature set performs optimally (i.e. contains a Markov boundary) is high, such as in Examaple 3, no feature set, produced by any feature selection algorithm or human expert, can perform statistically significantly better than random.*

**Remark 11: Performing Random Feature Selection Multiple Times.** *Theorem 1 defines how to compute $p = P(n; |\mathbf{V}|)$, the probability of one random feature selection of size $n$ resulting in a Markov blanket. Now we consider $k$ independent random feature selections, which correspond to a binomial distribution with parameters $k$ and $p$. The probability for finding at least one Markov blanket when conducting $k$ random selections of size $n$ is $1 - (1 - p)^k$.*

*For $p = 20\%$, a relatively low probability of obtaining a Markov blanket when conducting random feature selection only once. When we conduct random feature selection 10 times, the probability of obtaining at least one Markov blanket is fairly high, i.e. $1 - (1 - 20\%)^{10} \approx 89\%$.*

**Theorem 2: Monotonicity of $P(n; |\mathbf{V}|)$ w.r.t size of randomly selected feature set.** *In all distributions, as the size $n$ of the randomly selected feature set size increases, the probability of obtaining a Markov blanket, thus achieving optimal predictivity, increases monotonically. The probability stays unchanged from size $n$ to size $n + 1$ random selection only when either the probability is zero at $n$ and $n + 1$ (when $n + 1 < \min |\mathbf{MB}_i|$) or the probability has reached one at $n$.*

*Proof Sketch*
As we randomly select more features, both the number of Markov blankets and the number of all possible subsets to choose from would change. Here, we prove that the ratio of these two numbers, or probability of hitting a Markov blanket, increases monotonically. Moreover, the second part of the theorem states that when probability is not zero or one, it *strictly increases* monotonically.

To prove the first part of the theorem, one needs to look at how the numerator (size of all Markov blankets of size $n$ or $|\mathcal{MBL}^n|$) and denominator (size of all subsets of size $n$ or $|\mathcal{P}^n|$) are changing with $n$. It is not difficult to show the denominator changes by factor $(|\mathbf{V}| - n)/(n+1)$ going from $n$ to $n+1$. Then, we need to show that $|\mathcal{MBL}^{n+1}|$ increases at least with this factor compared to $|\mathcal{MBL}^n|$. This can be done by counting the subset relations from $\mathcal{MBL}^n$ to $\mathcal{MBL}^{n+1}$. This can be broken down in to two situations. First, if only $\mathbf{MBL} \in \mathcal{MBL}^n$ and no sets of size $n$ are subsets of $\forall \mathbf{MBL} \in \mathcal{MBL}^{n+1}$, the numerator also changes with $(|\mathbf{V}| - n)/(n+1)$. Second, if $\mathbf{MBL} \in \mathcal{MBL}^{\mathbf{n}}$ and some other sets of size $n$ are subsets of at least one $\mathbf{MBL} \in \mathcal{MBL}^{n+1}$, the ratio of change in the numerator when going from $n$ to $n+1$ is larger than $(|\mathbf{V}| - n)/(n+1)$.

The proof for the second part of theorem corresponds to the first case above, i.e. when sets in $\mathcal{MBL}^{n+1}$ do not have any subset of size $n$ that is not in $\mathcal{MBL}^n$. We will show that this results in either $\mathcal{MBL}^n = \mathcal{MBL}^{n+1} = \varnothing$ or $\mathcal{MBL}^n = \mathcal{P}^n$, i.e. probabilities 0 or 1. The argument comes down to the fact that $\mathcal{P}^n$ can be traversed with single variable exchange and one cannot partition it to two parts $\mathcal{MBL}^n$ and $\mathcal{P}^n \backslash \mathcal{MBL}^n$ where the distance between any pair of sets has to be $\geq 2$.

*Proof of Theorem 2.* Consider the set of all subsets or *power set* of $\mathbf{V}$, denoted as $\mathcal{P} = \{\mathbf{P} | \mathbf{P} \subseteq \mathbf{V}\}$. $\mathcal{P}^n$ is a set of subsets of $\mathbf{V}$ with size $n$. To prove the monotonicity of the probability of obtaining a Markov blanket with respect to the randomly selected feature set size, is to prove

$$P(n; |\mathbf{V}|) = \frac{|\mathcal{MB}^n|}{|\mathcal{P}^n|} \leq \frac{|\mathcal{MBL}^{n+1}|}{|\mathcal{P}^{n+1}|} = P(n+1; |\mathbf{V}|) \tag{30}$$

The ratio of $\mathcal{P}^{n+1}$ to $\mathcal{P}^n$ is easy to compute,

$$\frac{|\mathcal{P}^{n+1}|}{|\mathcal{P}^n|} = \frac{\binom{|\mathbf{V}|}{n+1}}{\binom{|\mathbf{V}|}{n}} = \frac{|\mathbf{V}| - n}{n+1}. \tag{31}$$

Above ratio is visualized in the Hasse diagram in Fig. 1. This diagram represents the *partially ordered set* of the power set $\mathcal{P}$ with the *partial order* being *proper subset* $\subset$ relation (also denoted as $\subsetneq$). To be more contained and clear we explain this diagram in more details below.

In this Hasse diagram, the nodes represent subsets of $\mathbf{V}$, arranged in layers by subset size, i.e. $|\mathbf{X}| = n$ if node $\mathbf{X}$ is in layer $n$. The edges represent inclusion ordering $(\subset)$ for nodes in adjacent layers. These edges are implicitly directed from top to bottom. If a node representing a set $\mathbf{X}$ in layer $n$ is connected by an edge to a node representing a set $\mathbf{Y}$ in layer $n+1$, we have $\mathbf{X} \subset \mathbf{Y}$.
We first illustrate obtaining $\frac{|\mathcal{P}^{n+1}|}{|\mathcal{P}^n|}$ through the Hasse Diagram. Each node representing a subset of $V$ with size $n$, i.e. $\forall \mathbf{P} \in \mathcal{P}^n$, has $|\mathbf{V}| - n$ out-going edges, since any subset of size $n$ can be padded into a subset with size $n+1$ by adding one additional variable that is not currently in the subset. Each node representing a subset of $V$ with size $n+1$, i.e. $\forall \mathbf{P} \in \mathcal{P}^{n+1}$, has $n+1$ incoming edges, since each subset with size $n+1$ can be obtained by adding one element to a subset with size $n$, with each element in the subset of size $n+1$ as a possible addition. Given the total number of outgoing edges from $\mathcal{P}^n$ must equal the total number of incoming edges to $\mathcal{P}^{n+1}$, i.e. $|\mathcal{P}^n|(|\mathbf{V}| - n) = |\mathcal{P}^{n+1}|(n+1)$, resulting in Equation (31).

We next consider $\frac{|\mathcal{MBL}^{n+1}|}{|\mathcal{MBL}^n|}$. A subset of Markov blanket of size $n$ is a member of $\mathcal{P}^n$. Therefore, the Markov blanket sets has all the properties discussed above. A Markov blanket of size $n$ has $|\mathbf{V}| - n$ out going edges, each connecting to a Markov blanket of size $n+1$. A Markov blanket of size $n+1$ has $n+1$ incoming edges, however, not all $n+1$ edges need to come from a Markov blanket of size $n$. For example, consider $\mathcal{MBL}^2 = \{\{A_1, B_1\}, \{A_1, B_2\}, \{A_2, B_1\}, \{A_2, B_2\}\}$. The Markov blanket of size 3 $\{A_1, A_2, B_1\}$ has 3 incoming edges, two from members of $\mathcal{MBL}^2$, i.e. $\{A_1, B_1\}$ and $\{A_2, B_1\}$, and one from set $\{A_1, A_2\}$. Let $q$ be the number of incoming edges into a Markov blanket of size $n+1$ from Markov blankets of size $n$, $0 \leq q \leq n+1$. Note that when $q = 0$, the corresponding Markov blanket of size $n+1$ is a Markov boundary, since none of its subsets of size $n$ is a Markov blanket.

Given the number of incoming and outgoing edges must be equal, we have,

$$|\mathcal{MBL}^n|(|\mathbf{V}| - n) = \sum_{i=1}^{|\mathcal{MBL}^{n+1}|} q_i \tag{32}$$

$$\leq |\mathcal{MBL}^{n+1}|(n+1), \tag{33}$$

where $q_i$ is the number of $\mathcal{MBL}^n$ sets that are included in $i$-th member of $\mathcal{MBL}^{n+1}$. Putting together Equation (31) and Equation (33), we obtain Equation (30), completing the monotonicity proof.

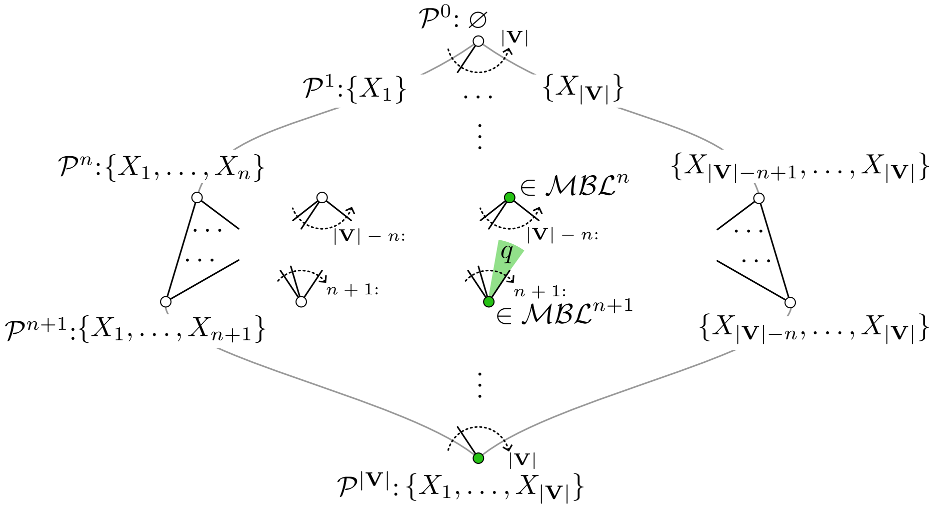

Figure 1: The figure shows the Hasse diagram of $\mathcal{P}$, i.e. the power set of $\mathbf{V}$, in a partial order by the inclusion ($\subset$) relation. The nodes in the diagram represents individual subsets arranged in layers by subset size: layer 0 (at the top) contains the empty set, and each subsequent lower layer contains subsets of increasing sizes. The edges in the diagram represents inclusion ($\subset$) relationships for nodes in adjacent layers. If a node representing a set $\mathbf{X}$ in layer $n$ is connected to a node representing a set $\mathbf{Y}$ in layer $n+1$, then $\mathbf{X} \subset \mathbf{Y}$. The subsets or nodes that are Markov blankets are shown with filled circles. The number of incoming edges to a Markov blanket of size $n+1$ from a Markov blanket of size $n$ is denoted as $q$, $0 \leq q \leq n+1$.

Next we show that the equal sign in Equation (30) can only corresponds to either $P(n; |\mathbf{V}|) = P(n+1; |\mathbf{V}|) = 0$ or $P(n; |\mathbf{V}|) = P(n+1; |\mathbf{V}|) = 1$.

To get equality in Equation (30), we need to have $\forall i \in \mathcal{MBL}^{n+1} : q_i = n+1$ in Equation (33). This, as we will show below, results in either $\mathcal{MBL}^n = \mathcal{P}^n$, i.e. all subsets of size $n$ are Markov blankets, thus $P(n; |\mathbf{V}|) = 1$, or $\mathcal{MBL}^n = \mathcal{MBL}^{n+1} = \varnothing$, thus $P(n; |\mathbf{V}|) = P(n+1; |\mathbf{V}|) = 0$.

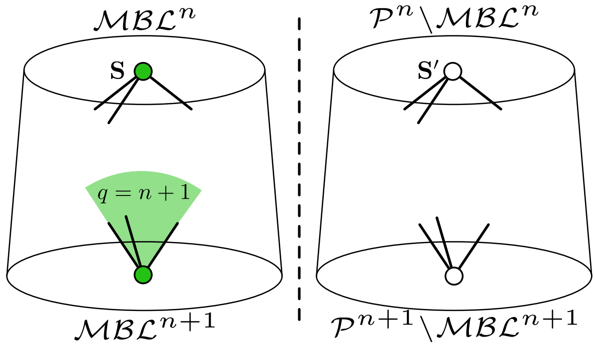

Figure 2: When $\forall i \in \mathcal{MBL}^{n+1} : q_i = n + 1$, no set in $\mathcal{MBL}^{n+1}$ includes any set $\mathbf{S}' \in \mathcal{P}^n \backslash \mathcal{MBL}^n$. It follows that $\forall \mathbf{S} \in \mathcal{MBL}^n, \forall \mathbf{S}' \in \mathcal{P}^n \backslash \mathcal{MBL}^n$ we must have $|\mathbf{S}'$
$\mathbf{S}| \geq 2$ otherwise the one extra member in $\mathbf{S}'$ can be added to $\mathbf{S}$ to make a set $\mathbf{S}_1 := \mathbf{S} \cup \mathbf{S}' \in \mathcal{MBL}^{n+1}$ which connects $\mathbf{S}'$ to $\mathcal{MBL}^{n+1}$ and this can not happen when we have all $q_i = n + 1$. On the other hand, $\mathcal{P}^n$ contains all the subsets of size $n$ and can be traversed with a single variable exchange operation, i.e. there is no way of partitioning $\mathcal{P}^n$ to two with all distances between members of these partitions $\geq 2$.

In Equation (33), when all $q_i = n + 1$, all members of $\mathcal{MBL}^{n+1}$ only includes members of $\mathcal{MBL}^n$ and they have no connections to subsets that are not Markov blankets of size $n$, i.e. $\mathcal{P}^n \backslash \mathcal{MBL}^n$. This is visualized in in Fig. 2. This means any set $\mathbf{S} \in \mathcal{MBL}^n$ and set $\mathbf{S}' \in \mathcal{P}^n \backslash \mathcal{MBL}^n$ must have at least two variables difference between them ($|\mathbf{S} \backslash \mathbf{S}'| \geq 2$ and $|\mathbf{S}' \backslash \mathbf{S}| \geq 2$). This is because when there is only one variable difference, $\mathbf{S}' \in \mathcal{P}^n \backslash \mathcal{MBL}^n$ will get included in a set in $\mathcal{MBL}^{n+1}$ by adding the variable $X = \mathbf{S} \backslash \mathbf{S}'$ to $\mathbf{S}'$, creating a connection between $\mathcal{P}^n \backslash \mathcal{MBL}^n$ and $\mathcal{MBL}^{n+1}$, which is not allowed when we have to set all existing $q_i = n + 1$.

However, having variables difference between all possible $S \in \mathcal{MBL}^n$ and $S' \in \mathcal{P}^n \backslash \mathcal{MBL}^n$ is not possible. Because there is always a sequence of subsets of size $n$ with one variable exchange between $\mathbf{S}$ and $\mathbf{S}'$. And since $\mathbf{S} \in \mathcal{MBL}^n$ and $\mathbf{S}' \in P^n \backslash \mathcal{MBL}^n$ and $\mathcal{MBL}^n \cup \mathcal{P}^n \backslash \mathcal{MBL}^n = \mathcal{P}^n$, there has to exist two neighboring sets in this sequence where one set is in $\mathcal{MBL}^n$ and the other in $\mathcal{P}^n \backslash \mathcal{MBL}^n$. This creates a contradiction, since these neighboring sets has distance of 1.

There are only two ways to resolve this contradiction: to have $\mathcal{MBL}^n = \mathcal{MBL}^{n+1} = \varnothing$ or to have $\mathcal{MBL}^n = \mathcal{P}^n$. The former case corresponds to $P(n; |\mathbf{V}|) = P(n + 1; |\mathbf{V}|) = 0$, which happens when the randomly selected feature set size is smaller than the size of the smallest Markov boundary, i.e. $n + 1 < \min |\mathbf{MB}_i|$. In this case, there is no way to select a subset that is a Markov blanket. Therefore, we have $|\mathcal{MBL}^n| = |\mathcal{MBL}^{n+1}| = 0$, and we will get equality in Equation (30). The latter case corresponds to $P(n; |\mathbf{V}|) = 1$, which happens when all subsets of size $n$ and thus all subsets of larger sizes are Markov blankets. This completes the proof.

□

**Example 4.** *Consider a small example,*

$$\mathbf{V} = \{A, B, C, D\}, \tag{34}$$

$$\mathbf{MB} = \{\{A\}, \{C, D\}\}. \tag{35}$$

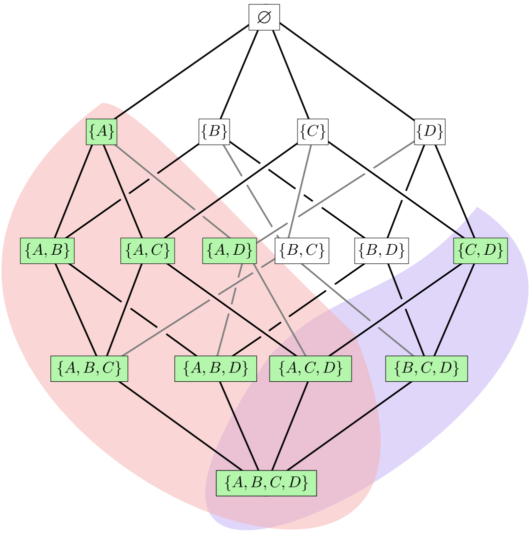

Figure 3: The Hasse diagram representing power set $\mathcal{P}$ for Example 4. Nodes representing Markov blankets are colored in green. Red shading and purple shading represent the Markov blankets corresponding to Markov boundaries $\{A\}$ and $\{C, D\}$, respectively. The probability of randomly selecting a subset of size $n = 0, 1, 2, 3, 4$ that includes a Markov boundary is given by $0, \frac{1}{4}, \frac{2}{3}, 1, 1$, respectively.

*The Hasse diagram for the power set of* $\mathbf{V}$ *is shown in Fig. 3 and we have* $P(0) = 0, P(1) = 1/4, P(2) = 2/3, P(3) = 1, P(4) = 1$.

**Theorem 3: p-value of Markov boundary or the Markov boundary feature selector in a faithful distribution.**

a) *The unique Markov boundary* $\mathbf{MB}$ *of* $T$ *(or a feature selector* $F_{\mathbf{MB}}$ *that selects the Markov boundary) has the unique smallest p-value and thus is the maximally statistically distinguishable from random among all variable sets with the same or larger sizes (or feature selectors that selects those sets), and among all variable sets with optimal performance (or feature selectors that selects those sets). If significance level (alpha) is chosen to be feasible relative to the number of unique random sets (*$\alpha^{-1} \leq$ *the number of unique random sets with size* $|\mathbf{MB}|$*) then the p-value* $\mathbf{MB}$ *will be significant.*

b) *The p-value of the Markov boundary (or the feature selector that selects the Markov boundary can be directly calculated from the total number of variables and the size of the unique Markov boundary,*

$$p = 1 / \binom{|\mathbf{V}|}{|\mathbf{MB}|}. \tag{36}$$

*Proof.* Note that the p value of a feature set and the p-value of the feature selector that produces the feature set on a particular dataset can be described with the same underlying statistical properties. In the proof, we will focus on the feature set, but the proof extends to the feature selector that produces the feature set.

i) We first prove the unique Markov boundary $\mathbf{MB}$ has the smallest p-value among all feature sets of the same size, i.e $|\mathbf{MB}|$.

The Markov boundary contain all necessary information regarding $T$, and results in optimal performance for $T$ (Definition 1 and Remark 1). Consider Definition 7, for feature set of the same size as $\mathbf{MB}$, the set $\mathbf{S}_{\mathrm{F}} = \mathbf{MB}$ maximizes the performance metric $M(\mathbf{S}_{\mathrm{F}}, T, D)$ in the large sample. Therefore, $p$ is minimized for $\mathbf{MB}$, resulting in maximal statistical distinguishability.

ii) Next, we prove that for feature set $\mathbf{S}_\mathrm{F}$ where $|\mathbf{S}_\mathrm{F}| > |\mathbf{MB}|$, the Markov blankets of size $|\mathbf{S}_\mathrm{F}|$ has the smallest p-value.

A Markov blanket contains all information regarding $T$ and results in optimal performance given a sufficiently regularized classifier (Definition 2 and Remark 2). Therefore, when $|\mathbf{S}_\mathrm{F}| > |\mathbf{MB}|$, $\mathbf{S}_\mathrm{F}$ where $\mathbf{S}_\mathrm{F} \supset \mathbf{MB}$ (i.e. $\mathbf{S}_\mathrm{F}$ is a Markov Blanket) maximizes $M(\mathbf{S}_\mathrm{F}, T, D)$ over all feature sets of size $|\mathbf{S}_\mathrm{F}|$. Therefore, given Definition 7, p-value is minimized when $\mathbf{S}_\mathrm{F}$ is a Markov blanket of $T$ over all feature sets of size $|\mathbf{S}_\mathrm{F}|$, where $|\mathbf{S}_\mathrm{F}| > |\mathbf{MB}|$.

iii) Given the monotonicity Theorem 2, the $p$-value for the Markov boundary is smaller than any Markov blanket of a larger size. Putting this together with i) and ii), we establish that the Markov boundary has the smallest $p$ value among all variable sets with the same or larger sizes and among all variable sets with optimal performance (i.e. Markov blankets).

iv) Given the Definition 7 of the better-than-random-test, it is easy to see that the p-values of the Markov boundary or Markov blankets of $T$ from the better-than-random test is equivalent to the probability of obtaining a Markov boundary or Markov blanket as stated in theorem 1, and can be computed with equations (6) through (8).

For faithful distribution with one unique Markov boundary, the p-value of the Markov boundary or the probability of obtaining a Markov boundary when sampling variable sets of size $|\mathbf{MB}|$ is:

$$p = P(n = |\mathbf{MB}|; \mathbf{V}) = 1 / \binom{|\mathbf{V}|}{|\mathbf{MB}|}, \tag{37}$$

which is the simplest case of equation (6) because the Markov boundary is unique. This $p$ value will be significant at $\alpha = 0.05$ when $\binom{|\mathbf{V}|}{|\mathbf{MB}(T)|} > 20$, which includes most real world distributions where $|\mathbf{V}| \gg |\mathbf{MB}|$ and $|\mathbf{MB}| \sim 10^1$.

For faithful distribution with one unique Markov boundary, the p-value of the Markov blankets $\mathbf{MBL}$ or the probability of obtaining a Markov blanket when sampling variable sets of size $|\mathbf{MBL}|$ is:

$$p = P(n = |\mathbf{MBL}|; \mathbf{V}) = \binom{|\mathbf{V}| - |\mathbf{MB}|}{|\mathbf{MBL}| - |\mathbf{MB}|} \Big/ \binom{|\mathbf{V}|}{|\mathbf{MBL}|}. \tag{38}$$

To summarize, we proved: For Theorem 3a and 3b, the Markov boundary has the smallest p-value among all sets with the same size (part i), all sets with optimal performance (part ii), and all variable sets of larger sizes (part iv and part ii). And for Theorem 3c, formula for computing the p-value of $\mathbf{MB}$ are given (part iii). □

**Remark 12: p-value of MB is significant in most faithful distributions.** *For most real world distributions, where $|\mathbf{V}| \gg |\mathbf{MB}|$ and $|\mathbf{MB}| \sim 10^1$, p-value of $\mathbf{MB}$ would be significant considering standard $\alpha$ levels. For example, $p \approx 2 \times 10^{-4}$ for $|\mathbf{V}| = 30$ and $|\mathbf{MB}| = 3$ and usually we have more variables on both sets , resulting in smaller p-values.*

**Theorem 4: p-value of Markov boundary or the Markov boundary feature selector in MBC-TIE distributions.**

a) *The Markov boundary with the smallest size, denoted as $\mathbf{MB}_s$, (or a feature selector that selects the Markov boundary) has the smallest p-values among all feature sets of the same size or larger (or feature selectors that selects those sets), and among all sets with optimal performance (or feature selectors that selects those sets).*

*If the smallest $\mathbf{MB}$ is unique, then its p-value will be significant when $\alpha$ is chosen to be feasible relative to number of random sets ($\alpha^{-1} \leq$ the number of unique random sets with size $|\mathbf{MB}_s|$).*

b) *An arbitrary member of $\mathcal{MB}$ (or a feature selector that selects that set) is not guaranteed to have smallest p-value among all feature sets with optimal performance. The p-value is not guaranteed to be unique, nor statistically significant under the similar conditions to part a. However, it is guaranteed to have smallest p-value among all feature sets with the same size or larger.*

*Proof.*

a) Proof of part a are similar to the proof of Theorem 3a and 3b. Markov boundaries and Markov blankets of any sizes have the smallest p-value among all sets of the same size, due to optimal performance. Further, sets with optimal performance are Markov boundaries or Markov blankets, i.e. members of $\mathcal{MBL}$. Due to the monotonicity Theorem 2, any member $\mathbf{MBL} \in \mathcal{MBL}$ would have p-values greater than or equal to the p-value of $\mathbf{MB}_s$.

   The formula for computing the p-value for the unique smallest $\mathbf{MB}$ is the same as equation (37). Its p-value will be significant when alpha is chosen to be feasible relative to the number of random sets ($\alpha^{-1} \leq$ the number of unique random sets with size $|\mathbf{MB}_s|$).

b) The p-value of a arbitrary member of $\mathbf{MB}$ is not guaranteed to be smallest, as proved in part i. It is also not guaranteed to be unique since Markov boundaries and Markov blankets of the same sizes would result in the same p-value. This p-value can be computed using equations (6) through (8). The p-value is not guaranteed to be significant, since the proportion of sets with optimal performance (i.e. Markov boundaries and Markov blankets of the same size) over the number of all possible sets at that sample size could be larger than standard $\alpha$. In Example 2, for variable set size of 2, we have 6 Markov boundaries among 28 possible random sets of size 2, which results in $p = 3/14$, greater than standard $\alpha = 0.05$. Or in Example 4, considering $\mathbf{MB}_2 = \{C, D\}$, it's p-value is $p = 2/3$. This value is very large because of all the Markov blankets resulted from $\mathbf{MB}_1 = \{A\}$.

   An arbitrary member of $\mathcal{MB}$ is guaranteed though to have the smallest p-value among all sets of the same size due to its optimal performance. In Example 2, $p = 3/14 > 0.05$, but this is the smallest p-value possible for $n \geq 2$.

□

**Corollary 4.1: Impossibility of better-than-random.** *In infinitely many distributions, it is impossible for any algorithm to select a feature set with optimal predictivity that is statistically better than random feature selection of the same size.*

*Proof.* Following Theorem 4a, $\mathbf{MB}_s$ has the smallest p-values among all feature sets with optimal performance. In distributions where the number of Markov boundaries of size $|\mathbf{MB}_s|$ over the number of subsets of size $|\mathbf{MB}_s|$ is larger than $\alpha$, no feature set of size $|\mathbf{MB}_s|$ can achieve statistical significance in the better-than-random test. Given Theorem 2 any feature set of size larger than $|\mathbf{MB}_s|$ also can not achieve statistical significance in the better-than-random test. Therefore, it is impossible for any algorithm to select a feature set with optimal predictive performance that is statistically better-than random in these distributions. □

**Theorem 5: Causal significance of feature set with smallest p-value.** *If we do not impose restrictions on the data distributions, statistical distinguishability from random sets cannot determine the causal significance of selected feature sets.*

a) *In faithful distributions under optimal predictivity, the causal feature set has the smallest p-value compared to all feature sets of the same size or larger.*

b) *In faithful distributions under suboptimal predictivity, causal feature sets may not be significant while non-causal feature sets achieve statistical significance according to the better-than-random test.*

c) *In MBC-TIE distributions, regardless of predictivity, the causal feature set is not guaranteed to have the smallest p-value or achieve statistical significance.*

*Proof.*

a) In faithful distributions, **MB** corresponds to the direct causes, direct effects, and direct causes of direct effects of $T$ and it has optimal performance (Remark 1). Also, Theorem 3a showed that **MB** has the smallest p-value among all feature sets with optimal performance. Therefore, in faithful distribution and under optimal performance, the causal feature set has the smallest p-value compared to all feature sets of the same size or larger.

b) In faithful distributions, we are at a feature set size that is smaller than the size of the Markov boundary, optimal predictivity can not be reached. We gave an example below to show that under this condition, a non-causal feature set can have smaller p-value compared to a causal feature set. And a non-causal feature set can have significant p-value while a causal feature set has a non-significant p-value.

**Example 5.** *Let $\langle \mathbf{V}, G, P \rangle$ denote a Causal Probability Network with the structure in Figure 4. $P$ denotes a distribution faithful to $G$. The target of interest is $T$. The Markov boundary of $T$ is $\{X_1, X_2\}$. Variables $I_i$, where $i = \{1, 2, \ldots, 22\}$ do not contain information regarding $T$. $M(\mathbf{S}, T, D) = \text{maximize}_{\mathbf{C}} M(\mathbf{S}, \mathbf{C}(T, D \downarrow \mathbf{S}))$ denotes the value of $M$ maximized over the set of classifier learners $\mathbf{C}$ for predicting $T$ using feature set $\mathbf{S} \subseteq \mathbf{V}$ and data sample $D$.*

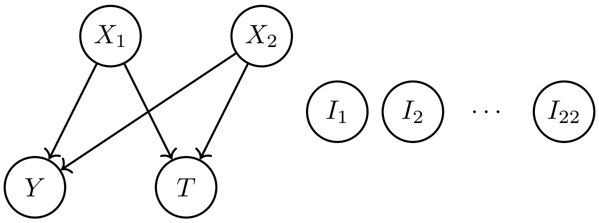

Figure 4: Graph $G$, specifying the faithful Causal Probabilistic Network $\langle \mathbf{V}, G, P \rangle$.

*This graph admits an infinite number of parameterizations such that,*

$$\forall i \in \{1, 2\}, \forall j \in \{1, 2, \ldots, 22\} : M(\{Y\}, T, D) > M(\{X_i\}, T, D) > M(\{I_j\}, T, D). \tag{39}$$

*In other words, for these parameterizations, $\{Y\}$ contains more information regarding the target compared to any other feature set of size 1. As a result, when selecting a variable set of size one, the feature set that is the most distinguishable from random would be set $\{Y\}$, not a set containing the causes of $T$.*

*We illustrate this with one specific parameterization where all variables are binary:*

$$\begin{aligned}
&P(X_1 = 1) = 0.4 \\
&P(X_2 = 1) = 0.6 \\
&P(I_i = 1) = 0.5 \\
&P(T = 1|X_1 = 1, X_2 = 1) = 0.1 \\
&P(T = 1|X_1 = 1, X_2 = 0) = 0.9 \\
&P(T = 1|X_1 = 0, X_2 = 1) = 0.9 \\
&P(T = 1|X_1 = 0, X_2 = 0) = 0.1 \\
&P(Y = 1|X_1 = 1, X_2 = 1) = 0.9 \\
&P(Y = 1|X_1 = 1, X_2 = 0) = 0.1 \\
&P(Y = 1|X_1 = 0, X_2 = 1) = 0.1 \\
&P(Y = 1|X_1 = 0, X_2 = 0) = 0.9
\end{aligned} \tag{40}$$

*We use opposite of 0-1 loss as metric $M(\mathbf{S}, T, D)$ and assume infinite sample and best possible classification performance, i.e. $M(\mathbf{S}, T, D) = 1 - \mathbb{E}_{\mathbf{S}}\left[\min(P(T = 1|\mathbf{S}), 1 - P(T = 1|\mathbf{S}))\right]$, where $\mathbf{S}$ denote a feature set. $\mathbb{E}_{\mathbf{S}}\left[\min(P(T = 1|\mathbf{S}), 1 - P(T = 1|\mathbf{S}))\right]$ is the optimal Bayes error for predicting $T$ using set $\mathbf{S}$.*

*Given the parameterization in equation set (40), $M(\{Y\}, T, D) = 0.82 > M(\{X_i\}, T, D) = 0.58 > M(\{I_j\}, T, D) = 0.516$, for any $i \in \{1, 2\}$ and $j \in \{1, 2, \ldots, 22\}$.*

*For feature set $\{Y\}$, the corresponding random feature selection distribution comprises $\binom{25}{1} = 25$ predictive performances from all feature sets of size 1. Applying Definition 7, the p-value of $\{Y\}$, is*

$$p = 1 - \sum_{M < M(\{Y\}, T, D)} P_1(M(\mathbf{S}^1, T, D)) = 1 - \frac{24}{25} = 0.04. \tag{41}$$

*The p-value of $\{X_i\}$ , for any $i \in \{1, 2\}$, is*

$$p = 1 - \sum_{M < M(\{X_i\}, T, D)} P_1(M(\mathbf{S}^1, T, D)) = 1 - \frac{22}{25} = 0.12. \tag{42}$$

*The p-value of $\{I_j\}$, for any $j \in \{1, 2, \ldots 22\}$, is*

$$p = 1 - \sum_{M < M(\{I_j\}, T, D)} P_1(M(\mathbf{S}^1, T, D)) = 1 - \frac{0}{25} = 1. \tag{43}$$

*Since $M(\{Y\}, T, D)$ has the largest value among all feature sets of size 1, $\{Y\}$ has the smallest p-value among all feature sets of size 1. However, $Y$ is not a cause of $T$.*

*This example illustrates that when the selected feature set size is smaller than that of the* **MB**, *classes of faithful distributions exist where feature sets containing causal features of $T$ (e.g. direct causes, direct effects, direct causes of direct effects) have larger p-values than those that do not contain causal features.*

c) In MBC-TIE distributions, the causal feature set is not guaranteed to have the smallest p-value or achieve statistical significance. This is regardless of whether we assume optimal predictivity is achievable or not. Under suboptimal predictivity, the proof is similar to what is outlined in part b. Even under optimal predictivity (i.e. among feature sets that are Markov boundaries or Markov blankets), the causal feature set is not guaranteed to have the smallest p-value. This is because the Markov boundary set that is the smallest in size is not necessarily causal. We illustrate this with an example.

**Example 6.** *We use the same causal network structure as Example 5 shown in Figure 4. Unlike Example 5, $P$ denotes a MBC-TIE distribution NOT faithful to $G$. The target of interest is $T$. There are multiple Markov boundaries of $T$, i.e. $\{Y\}$ and $\{X_1, X_2\}$. Variables $I_j$, where $j \in \{1, 2, \dots, 22\}$, do not contain information regarding $T$. A specific parameterization of $P$ is the following:*

$$P(X_1 = 1) = 0.4, P(X_2 = 1) = 0.6, P(I_j = 1) = 0.5, \tag{44}$$

*and,*

| $(X_1, X_2)$ | (1, 1) | (1, 0) | (0, 1) | (0, 0) |
|---|---|---|---|---|
| $P(T = 1 \mid X_1, X_2)$ | 0.1 | 0.9 | 0.9 | 0.1 |
| $P(Y = 1 \mid X_1, X_2)$ | 0.5 | 0 | 0 | 0 |
| $P(Y = 2 \mid X_1, X_2)$ | 0.5 | 0 | 0 | 0 |
| $P(Y = 3 \mid X_1, X_2)$ | 0 | 1 | 0 | 0 |
| $P(Y = 4 \mid X_1, X_2)$ | 0 | 0 | 1 | 0 |
| $P(Y = 5 \mid X_1, X_2)$ | 0 | 0 | 0 | 1 |

(45)

*Given the parameterization, the $M$ of feature sets of size 1 are:*

$$M(\{Y\}, T, D) = 0.9 > M(\{X_i\}, T, D) = 0.58 > M(\{I_j\}, T, D) = 0.516, \tag{46}$$

*for any $i \in \{1, 2\}$ and $j \in \{1, 2, \dots, 22\}$.*

*The $M$ values for feature sets of size 2 are:*

$$\begin{aligned} &M(\{X_i, Y\}, T, D) = M(\{X_1, X_2\}, T, D) = M(\{Y, I_j\}, T, D) = 0.9 \\ >&M(\{X_i, I_j\}, T, D) = 0.58 \\ >&M(\{I_j, I_k\}, T, D) = 0.516, \end{aligned} \tag{47}$$

*for any $i \in \{1, 2\}$ and $j, k \in \{1, 2, \dots, 22\}$ where $j \neq k$.*

*Given Definition 7, the variable set with the smallest p-value is $\{Y\}$, the smallest Markov boundary set:*

$$p = 1 - \sum_{M < M(Y,T,D)} P_1(M(\mathbf{S}^1, T, D)) = 1 - \frac{1}{25} = 0.04. \tag{48}$$

*The causal variable set $\{X_1, X_2\}$ has the same p-value as $\{X_i, Y\}$, and $\{I_j, Y\}$:*

$$p = 1 - \sum_{M < M(\{X_1, X_2\}, T, D)} P_2(M(\mathbf{S}^2, T, D)) = 1 - \frac{25}{300} = 0.0833. \tag{49}$$

*We can see that, the causal feature set has larger p-value compared to the non-causal Markov boundary set, and the p-value is not statistically significant at $\alpha = 0.05$.*

More interestingly, the p-value for the causal feature set can be arbitrarily close to one in a MBC-TIE distribution.

**Example 7.** *Consider the Causal Probabilistic Network $\langle \mathbf{V}, G, P \rangle$ with $G$ specified in 5. The multiple Markov boundaries are $\mathbf{MB} = \{\{X_i\} | i = 1, 2, \dots, m\}$. It is easy to see that as $m \to \infty$, $p \to 1$ for any Markov boundary.*

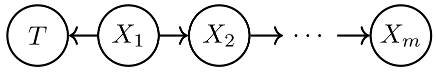

Figure 5: Graph $G$, specifying the structure of the Causal Probabilistic Network $\langle \mathbf{V}, G, P \rangle$.

□

### References

[1] Judea Pearl. *Probabilistic Reasoning in Intelligent Systems: Networks of Plausible Inference*. San Francisco, CA, USA: Morgan Kaufmann Publishers Inc., 1988. ISBN: 1558604790.

[2] Ioannis Tsamardinos and Constantin F Aliferis. "Towards principled feature selection: Relevancy, filters and wrappers". In: *International Workshop on Artificial Intelligence and Statistics*. PMLR. 2003, pp. 300–307.

[3] Constantin F Aliferis et al. "Local causal and Markov blanket induction for causal discovery and feature selection for classification part I: algorithms and empirical evaluation." In: *Journal of Machine Learning Research* 11.1 (2010).

[4] Alexander Statnikov et al. "Algorithms for discovery of multiple Markov boundaries". In: *Journal of Machine Learning Research* 14.Feb (2013), pp. 499–566.

[5] John Riordan. "An introduction to combinatorial analysis". In: (2014).
