## Supplementary material for "Optimal Predictivity of Random Signatures: Causes and Good Modeling Practices": Detailed_Methods

### Table of Contents

S1. Methods for Analytical Experiments on Microarray Data 1

S1.1 Datasets 1

S1.2 Comparing the Performance of Algorithmic Feature Selection vs. Random Feature Selection 2

S1.3 Number of Randomly Selected Features vs Predictive Performance 4

S1.4 Generalizability of Random Feature Selection 4

S2. Methods for Analytical Experiments on other Omics Data 5

S3. Methods for Empirical dissection of Markov Boundaries equivalence classes 5

#### S1. Methods for Analytical Experiments on Microarray Data

##### *S1.1 Datasets*

9 microarray datasets were used to (1) demonstrate the performance comparison between algorithmic feature selections vs. random feature selection with the same number of variables (Table 1), and (2) examine the change in performance of random feature selection as a function of the number of features selected (Figure 1a). In each datasets, the features are expression values of genes. Outcomes for all datasets are related to cancer diagnosis or prognosis.

Table S1. Characteristics of Microarray Datasets

| Data Name | # positive outcome | # obs | # features | Outcome | Reference |
|---|---|---|---|---|---|
| Bhattacharjee | 17 | 203 | 12600 | Lung caner vs Normal | 1 |
| Shipp | 19 | 77 | 5469 | Diffuse large B-cell lymphomas vs. folicular lymphamas | 2 |
| Singh | 50 | 102 | 12600 | Prostate tumor vs normal | 3 |
| MSKCC-PRCA | 13 | 92 | 26447 | Prostate tumor primary vs. metastic | 4 |
| Alon | 40 | 62 | 2000 | Colon tumors vs. normal | 5 |
| Neomark | 6 | 77 | 25702 | Oral Squamous cell carcinoma, recurrence vs. non-recurrence | 6 |
| Veer | 46 | 97 | 24188 | Breast Cancer 5 year Progression free survival | 7 |
| Metabric | 1328 | 1668 | 48803 | Breast Cancer 5 year survival | 8 |
| Ramaswamy | 12 | 76 | 13247 | Metastic vs. Primary Tumor | 9 |
| Iizuka* | 20 | 60 | 7070 | Hepatocellular carcinoma 1 year recurrence free survival | 10 |
| Yeoh* | 32 | 233 | 12240 | Acute lymphocytic leukemia relapse free survival | 11 |
| Rosenwald* | 113 | 227 | 7399 | Non-Hodgkin lymphoma 5 year survival | 12 |

| | | | | |
|---|---|---|---|---|
| Beer* | 24 | 86 | 7129 | Lung Adenocarcinoma 4 year survival |

* Experiments were conducted on these four datasets. Our results indicate that the predictive signal in these datasets are low (all algorithmic feature selection resulted in models with AUC<0.7). We excluded these datasets from our main analysis (table 1, figure 1a) since the goal of the study is to compare the algorithmic feature selection against random. This comparison is less meaningful when algorithmic feature selection results in low predictive performance. The performances on these datasets are shown in supplemental tableS5.

##### ***S1.2 Comparing the Performance of Algorithmic Feature Selection vs. Random Feature Selection***

The goal of this set of experiments is to compare the predictive performance of models with features selected by various Algorithmic feature selection methods vs. randomly selected feature set of the same size.

We used stratified repeated nested cross validation (NCV) for model selection and performance estimation with 10 folds with 1 repeat (none repeated performance estimator, NRPE) or 50 repeats (repeated performance estimator, RPE). The NCV procedure was run for each feature selection method and dataset combinations separately. The inner loop of the NCV is for model selection, where the combinations of hyperparameters for the classifier and the feature selection method that resulted in the best AUC was selected. The outter loop of the NCV estimate the predictive performance of the selected model. In the RPE setting, the NCV procedure was repeated 50 times to reduce biases associated with splitting the dataset into 10 folds. The average AUC from the outter loop was reported. The same data splits were used across individual experiments for different feature selection methods to allow direct comparison of predictive performances.

Algorithmic feature selection methods examined in this experiment were listed in Table S2. We chose these feature selection methods since they represent the families of feature selection methods that is often used for microarray data analysis. Hyperparameter for the feature selection methods were chosen following recommendations from prior benchmark studies (citation in Table S2).

We used polynomial kernel (SVM-poly) and logistic regression to build predictive models based on the selected features. Hyper-parameter selection for SVM-poly and selected feature selection methods are conducted using NCV. For SVM-poly, we considered the following hyper parameter combinations: C = {1e-3, 1e-2, 0.1, 1, 10, 100}, and d = {1, 2, 3, 4}. The SVM-poly results were reported in the main text of the paper (Table 1, Figure 1).

Table S2. Descriptions of feature selection methods.

| Method | Description |
|---|---|

| | |
|---|---|
| RFE[14] | SVM recursive feature elimination, with 50% of features eliminated at each iteration, as specified in the original publication[14]. Select feature set with best performance produced by SVM on the train test set. |
| GLL-OPT[15,16] | GLL-PC with hyperparameter selected from the Cartesian product of K={1,2,3}, alpha={0.01,0.05}. |
| RFVS[17] | Random Forest variables importance selection, with 20% elimination at each iteration rate. OOB error was used to select best feature set with 1 s.e. rule implemented. Feature importance was not recomputed at each iteration. This setting was chosen according to the original publication[17]. |
| relief[18,19] | relief feature selection with 1 neighbor, 50% of features eliminated at each iteration. Feature set selection with SVM backward wrapper. Hyperparameters chosen according to prior benchmark[15]. |
| LARS-EN[20] | Elastic net with alpha={0.25,0.5,0.75,1}, lambda={0.001, 0.01, 0.1, 1, 10, 100, 1000}; |
| UAF-KW[21,22] | Univariate feature ranking with Kruskal-Walis statistic with 20% reduction at each iteration. Feature selection with SVM backward wrapper . Hyperparameters chosen according to prior benchmark[15]. |
| SIMCA[23] | Soft Independent Modeling of Class Analogy: PCA based method where feature subset with best performance produced by SVM on the train test set were selected. |
| T-fdr | Select significant features according to fdr adjusted t test at alpha<0.05, according to what is commonly done as a first pass in differential expression analysis. |

To compare the predictive performance of random feature selection to that of an algorithmic feature selection, we generate an empirical distribution of the predictive performance of randomly selection feature sets with same number of features as the algorithmic feature selection in question. Specifically, for each algorithmic feature selection where we used NRPE or RPE 10 fold NCV for models selection and performance estimation, we run the same NCV procedure 100 times where features were random selected without replacement in all inner loop and outter loop with number of features matching that of the algorithmic feature selection. This results in 100 performance estimations from the random feature selection, establishing the empirical distribution of random feature selection. The NCV performance estimation of the algorithmic feature selection is compared to this empirical distribution to assess if the performance of the algorithmic feature selection is statistically significantly different from that of the random (Table 1, row 5 and 6) at $p<0.05$. We also compare the NCV performance estimation of the algorithmic feature selection to the mean of the 100 random feature selection performance estimations. This establish the added value of the algorithmic feature selection compared to random (Table 1, row 3).

##### *S1.3 Number of Randomly Selected Features vs Predictive Performance*

To examine the relationship between number of random features selected and the predictive performance, we ran another set of experiments where for each dataset, we randomly select 100 feature sets for each of the following sizes, {1, 2, 5, 10, 20, 50, 100, 200,

500, 1000, 2000, 5000, 10000, 20000, 50000}, without replacement. If the number of features in a dataset is smaller than a given random feature set size, that feature set size is not sampled. Model selection and performance estimation was conducted using 50 repeated RPE NCV. This experiment resulted in estimated predictive performance for each dataset and at different feature set sizes (Figure 1a).

##### *S1.4 Generalizability of Random Feature Selection*

To assess the ability of randomly selected features generalizing to an external dataset generated and assayed independently, we identified two pairs of datasets with overlapping gene expression features and clinical outcome. Models were built on one dataset (discovery data) of the pair and applied to the other dataset (validation) to assess generalizability, table S3 shows the source and characteristics of these datasets.

Table S3. Characteristics of datasets used to assess generalizability of random feature selection.

| **Discovery** | **Validation** | **Description** |
|---|---|---|
| Bhattacharjee[1]<br>Beer | Beer[13]<br>Bhattacharjee | Outcome of prediction is lung cancer vs normal. 7094 overlapping features. Bhattacharjee has 203 samples with 17 positive cases. Beer has 96 samples with 10 positive cases. |
| Bhattacharjee<br>Su | Su[24]<br>Bhattacharjee | Outcome of prediction is Adeno vs. squamous. 12533 overlapping features. Bhattacharjee has 160 samples with 21 positive cases. Su has 28 samples with 14 positive cases |

To evaluate performance of randomly selection feature sets on the discovery dataset, we implemented the same procedure in S1.3. To build the final model to be applied to the validation data, random feature selection was applied to the entire discovery dataset, and the SVM were built with the best hyperparameter selected in the NCV. The mean and variability of the performance of the 100 models built with randomly selected feature sets were reported. Results shown in Figure 1b.

We also produced results for the generalizability of algorithmic feature selection methods for comparison. Methods for assessing generalizability of algorithmic feature selection is similar to that of random feature selection, except each algorithmic feature selection produces one model to be evaluated in the validation data, whereas 100 models were produced for random feature selection to assess the variability in the randomly selected feature sets.

#### S2. Methods for Analytical Experiments on other Omics Data

To assess if the performance observed for random feature selection in microarray data holds in other omics data sets, we used datasets listed in Table S4. Performances for algorithmic feature selections were compared to random feature selection selecting feature set of the same size, as described in section S1.2. Results of these experiments were reported in Table 1, Figure 1a and supplemental table S7.

Table S4. Characteristics of Other Omics Datasets

| Data Name | Data Type | # positive outcome | # obs | # features | Outcome | Reference |
|---|---|---|---|---|---|---|
| MSKCC-miRNA | Copy Number | 13 | 92 | 368 | Prostate tumor primary vs. metastic | 4 |
| BRCA-Protein | Protein | 176 | 233 | 165 | Estrogen receptor negative vs. positive | 25,26 |
| BRCA-Methylation | Methylation | 176 | 233 | 27576 | Estrogen receptor negative vs. positive | 25,26 |
| Arcene | mass-spectromet ry | 398 | 900 | 10000 | Cancer (ovarian or prostate) vs. healthy | 27 |
| MacParland2018 | Signal Cell Gene Expression | 2621 | 3127 | 20007 | Hepatocytes vs other cell types | 28 |

#### S3. Methods for Empirical dissection of Markov Boundaries equivalence classes

Markov Boundary variables contain all information available about the target. According to our theoretical formula in Theorem 1, the probability of selecting a Markov bounary when randomly selection a fixed number of variables is related to the structure of Markov Boundaries and the total number of variables in a dataset. To illustrate this correspondence between random feature selection and the Markov Boundaries structure in real-world datasets. We applied the TIE* algorithm[29,30] to derive all Markov Boundaries of 2 microarray datasets Alon and Singh. We compared the probability of selecting at least one Markov boundaries for random feature selection with specific feature set sizes ({10,20 50,100:100:1000:1500:2000}) with the probability computed using our theoretical formula in Theorem 1c. The probability of selecting Markov boundaries using random feature selection were estimated emperically by random sampling feature sets of specific sizes 100 times. We estimated the theoretical probabilities according to Theorem 1c. The structure of the Markov Boundaries specified when applying Theorem 1c were approximated as the following. For the *m* Markov boundaries of a specific size *k*, find *k* integers $t_i$, where the product of $t_i$ approximates *m*. We start with $t_{i,i=1..k}$ =round($m^{1/k}$). The sumation of $t_i$ should also approximates the number of unique variables that constitutes the *m* Markov Boundaries. i.e. $t_{i,i=1..k}$ are members of the $N_{TG}$ set.This stratergy worked well for the Alon dataset. For the Sigh dataset, setting $t_{i,i=1..k}$ =round($m^{1/k}$) did not results in sumation of $t_i$ approximating the number of unique variables that constitutes the *m* Markov Boundaries. Upon closer inspection, we found many variables appeared in all Markov boundaries. Therefore, we set $t_{1,\ldots,j=}$1, where *j* is the number of variables that appeared in nearly all Markov boundaries (greater than the larger of 90% or the 70$^{th}$ percentile of the emperical distribution).And $t_{i,i=j..k}$ =round($m^{1/(k-j)}$). Being able to set $t_{1,\ldots,j}$=1 also made the application of the formula in Theorem 1c more computationally efficient for the Singh dataset, where each

Markov boundary has 29-38 members. Since after maniputation of Theorem 1c, $t_{1,\dots,j}$=1 does not need to appear in generating the power set of $N_{TG}$. In general, direct application of Theorem 1c as stated is computational infeasible for datasets with Markov boundaries with larger sizes. Further manipulation of Theorem 1c formula given the specific structures of Markov boundaries (as we did for Singh) or numerical approximations are needed. Results for Alon is shown in Figure 4c. Results for Singh in supplemental table S8b. After estimating the probability of selecting Markov boundaries of all sizes. We combine these estimates to estimate the probability of selecting at least one Markov boundary of any size. These probabilities were estimated for ramdomly selecting feature sets of specific sizes 1, 2, 5, and 10 times (Figure 4d).

We also mapped the variables discovered in the Markov boundaries of these datasets to gene pathways. Gene list was analyzed with Enrichr database[31], a database with the most comprehensive gene sets/libraries collections in the public domain. Enrichment analysis was done with methods used by Enrichr database including proportion test, Z-score and combined score which has shown better performance in benchmark comparisons[32] (PMID: 27141961).


1. Bhattacharjee, A., Richards, W.G., Staunton, J., Li, C., Monti, S., Vasa, P., Ladd, C., Beheshti, J., Bueno, R., Gillette, M., et al. (2001). Classification of human lung carcinomas by mRNA expression profiling reveals distinct adenocarcinoma subclasses. Proc. Natl. Acad. Sci. U.S.A. *98*, 13790–13795. https://doi.org/10.1073/pnas.191502998.

2. Shipp, M.A., Ross, K.N., Tamayo, P., Weng, A.P., Kutok, J.L., Aguiar, R.C.T., Gaasenbeek, M., Angelo, M., Reich, M., Pinkus, G.S., et al. (2002). Diffuse large B-cell lymphoma outcome prediction by gene-expression profiling and supervised machine learning. Nature Medicine *8*, 68–74.

3. Singh, D., Febbo, P.G., Ross, K., Jackson, D.G., Manola, J., Ladd, C., Tamayo, P., Renshaw, A.A., D'Amico, A.V., Richie, J.P., et al. (2002). Gene expression correlates of clinical prostate cancer behavior. Cancer cell *1*, 203–209.

4. Golub, T.R., Slonim, D.K., Tamayo, P., Huard, C., Gaasenbeek, M., Mesirov, J.P., Coller, H., Loh, M.L., Downing, J.R., Caligiuri, M.A., et al. (1999). Molecular classification of cancer: class discovery and class prediction by gene expression monitoring. Science *286*, 531–537. https://doi.org/10.1126/science.286.5439.531.

5. Alon, U., Barkai, N., Notterman, D.A., Gish, K., Ybarra, S., Mack, D., and Levine, A.J. (1999). Broad patterns of gene expression revealed by clustering analysis of tumor and

normal colon tissues probed by oligonucleotide arrays. Proc Natl Acad Sci U S A *96*, 6745–6750. https://doi.org/10.1073/pnas.96.12.6745.

6. Picone, M., Steger, S., Exarchos, K., De Fazio, M., Goletsis, Y., Fotiadis, D.I., Martinelli, E., and Ardigo, D. (2011). Enabling heterogeneous data integration and biomedical event prediction through ICT: the test case of cancer reoccurrence. Software Tools and Algorithms for Biological Systems, 367–375.

7. van 't Veer, L.J., Dai, H., van de Vijver, M.J., He, Y.D., Hart, A.A.M., Mao, M., Peterse, H.L., van der Kooy, K., Marton, M.J., Witteveen, A.T., et al. (2002). Gene expression profiling predicts clinical outcome of breast cancer. Nature *415*, 530–536. https://doi.org/10.1038/415530a.

8. Frishberg, A., Peshes-Yaloz, N., Cohn, O., Rosentul, D., Steuerman, Y., Valadarsky, L., Yankovitz, G., Mandelboim, M., Iraqi, F.A., Amit, I., et al. (2019). Cell composition analysis of bulk genomics using single-cell data. Nature Methods *16*, 327–332. https://doi.org/10.1038/s41592-019-0355-5.

9. Ramaswamy, S., Tamayo, P., Rifkin, R., Mukherjee, S., Yeang, C.H., Angelo, M., Ladd, C., Reich, M., Latulippe, E., Mesirov, J.P., et al. (2001). Multiclass cancer diagnosis using tumor gene expression signatures. Proc Natl Acad Sci U S A *98*, 15149–15154. https://doi.org/10.1073/pnas.211566398.

10. Iizuka, N., Oka, M., Yamada-Okabe, H., Mori, N., Tamesa, T., Okada, T., Takemoto, N., Sakamoto, K., Hamada, K., Ishitsuka, H., et al. (2005). Self-organizing-map-based molecular signature representing the development of hepatocellular carcinoma. FEBS Lett *579*, 1089–1100. https://doi.org/10.1016/j.febslet.2004.10.113.

11. Yeoh, E.-J., Ross, M.E., Shurtleff, S.A., Williams, W.K., Patel, D., Mahfouz, R., Behm, F.G., Raimondi, S.C., Relling, M.V., Patel, A., et al. (2002). Classification, subtype discovery, and prediction of outcome in pediatric acute lymphoblastic leukemia by gene expression profiling. Cancer Cell *1*, 133–143.

12. Rosenwald, A., Wright, G., Chan, W.C., Connors, J.M., Campo, E., Fisher, R.I., Gascoyne, R.D., Muller-Hermelink, H.K., Smeland, E.B., Giltnane, J.M., et al. (2002). The use of molecular profiling to predict survival after chemotherapy for diffuse large-B-cell lymphoma. N Engl J Med *346*, 1937–1947. https://doi.org/10.1056/NEJMoa012914.

13. Beer, D.G., Kardia, S.L., Huang, C.-C., Giordano, T.J., Levin, A.M., Misek, D.E., Lin, L., Chen, G., Gharib, T.G., Thomas, D.G., et al. (2002). Gene-expression profiles predict survival of patients with lung adenocarcinoma. Nature medicine *8*, 816.

14. Guyon, I., Weston, J., Barnhill, S., and Vapnik, V. (2002). Gene Selection for Cancer Classification using Support Vector Machines. Mach. Learn. *46*, 389–422. https://doi.org/10.1023/a:1012487302797.

15. Aliferis, C.F., Statnikov, A., Tsamardinos, I., Mani, S., and Koutsoukos, X.D. (2010). Local Causal and Markov Blanket Induction for Causal Discovery and Feature Selection for

Classification Part I: Algorithms and Empirical Evaluation. Journal of Machine Learning Research *11*, 171–234.

16. Aliferis, C.F., Statnikov, A., Tsamardinos, I., Mani, S., and Koutsoukos, X.D. (2010). Local Causal and Markov Blanket Induction for Causal Discovery and Feature Selection for Classification Part II: Analysis and Extensions. J. Mach. Learn. Res. *11*, 235–284.

17. Díaz-Uriarte, R., and Alvarez de Andrés, S. (2006). Gene selection and classification of microarray data using random forest. BMC Bioinformatics *7*, 3. https://doi.org/10.1186/1471-2105-7-3.

18. Kononenko, I. (1994). Estimating attributes: Analysis and extensions of RELIEF. In Machine Learning: ECML-94, F. Bergadano and L. De Raedt, eds. (Springer), pp. 171–182. https://doi.org/10.1007/3-540-57868-4_57.

19. Kira, K., and Rendell, L.A. (1992). A Practical Approach to Feature Selection. In Proceedings of the Ninth International Workshop on Machine Learning ML ’92. (Morgan Kaufmann Publishers Inc.), pp. 249–256.

20. Zou, H., and Hastie, T. (2005). Regularization and variable selection via the elastic net. Journal of the Royal Statistical Society: Series B (Statistical Methodology) *67*, 301–320. https://doi.org/10.1111/j.1467-9868.2005.00503.x.

21. Hollander, M., Wolfe, D.A., and Chicken, E. (2013). Nonparametric Statistical Methods (John Wiley & Sons).

22. Statnikov, A., Aliferis, C.F., Tsamardinos, I., Hardin, D., and Levy, S. (2005). A comprehensive evaluation of multicategory classification methods for microarray gene expression cancer diagnosis. Bioinformatics *21*, 631–643. https://doi.org/10.1093/bioinformatics/bti033.

23. WOLD, S., and SJÖSTRÖM, M. (1977). SIMCA: A Method for Analyzing Chemical Data in Terms of Similarity and Analogy. In Chemometrics: Theory and Application ACS Symposium Series. (AMERICAN CHEMICAL SOCIETY), pp. 243–282. https://doi.org/10.1021/bk-1977-0052.ch012.

24. Su, A.I., Welsh, J.B., Sapinoso, L.M., Kern, S.G., Dimitrov, P., Lapp, H., Schultz, P.G., Powell, S.M., Moskaluk, C.A., Frierson, H.F., et al. (2001). Molecular classification of human carcinomas by use of gene expression signatures. Cancer Res. *61*, 7388–7393.

25. The Cancer Genome Atlas Program (TCGA) - NCI (2022). https://www.cancer.gov/ccg/research/genome-sequencing/tcga.

26. Ray, B., Henaff, M., Ma, S., Efstathiadis, E., Peskin, E.R., Picone, M., Poli, T., Aliferis, C.F., and Statnikov, A. (2014). Information content and analysis methods for multi-modal high-throughput biomedical data. Sci Rep *4*, 4411. https://doi.org/10.1038/srep04411.

27. UCI machine learning repository Arcene Dataset. https://archive.ics.uci.edu/dataset/167/arcene.

28. MacParland, S.A., Liu, J.C., Ma, X.-Z., Innes, B.T., Bartczak, A.M., Gage, B.K., Manuel, J., Khuu, N., Echeverri, J., Linares, I., et al. (2018). Single cell RNA sequencing of human liver reveals distinct intrahepatic macrophage populations. Nature Communications *9*, 4383. https://doi.org/10.1038/s41467-018-06318-7.

29. Statnikov, A., and Aliferis, C.F. (2010). Analysis and Computational Dissection of Molecular Signature Multiplicity. PLOS Computational Biology *6*, e1000790. https://doi.org/10.1371/journal.pcbi.1000790.

30. Statnikov, A., Lytkin, N.I., Lemeire, J., and Aliferis, C.F. (2013). Algorithms for Discovery of Multiple Markov Boundaries. J Mach Learn Res *14*, 499–566.

31. Chen, E.Y., Tan, C.M., Kou, Y., Duan, Q., Wang, Z., Meirelles, G.V., Clark, N.R., and Ma'ayan, A. (2013). Enrichr: interactive and collaborative HTML5 gene list enrichment analysis tool. BMC Bioinformatics *14*, 128. https://doi.org/10.1186/1471-2105-14-128.

32. Kuleshov, M.V., Jones, M.R., Rouillard, A.D., Fernandez, N.F., Duan, Q., Wang, Z., Koplev, S., Jenkins, S.L., Jagodnik, K.M., Lachmann, A., et al. (2016). Enrichr: a comprehensive gene set enrichment analysis web server 2016 update. Nucleic Acids Res *44*, W90-97. https://doi.org/10.1093/nar/gkw377.
