## SupplementaryDocument2 for "Optimal Predictivity of Random Signatures: Causes and Good Modeling Practices"

**Uses of molecular signatures**
Our work is motivated by research projects in longevity/aging and cancer. In the domain of cellular senescence, SenMayo[1] is a signature that aims to identify senescent cells across tissues and species with high fidelity. Using this senescence panel, the authors were able to characterize senescent cells at the single cell level and identify key intercellular signaling pathways. Sensig[2] is an in vivo-derived senescence signature using a foreign body response-driven fibrosis model in a p16-CreERT2;Ai14 reporter mouse. As a third example, a signature comprising 68 differentially expressed RNAs that helps to discriminate between senescent and non-senescent cells was reported in[3] (also discussed in Li et al. [4]). In Wentcher et al.[3], signatures are used as part of the effort to identify different senescence programs that developed divergently, one in which cells expressed traditional senescence markers such as p16 (CDKN2A) mRNA, and another in which cells expressed long noncoding RNAs and splicing was dysregulated and to obtain evidence that the proliferation status at the time of senescence initiation affects the path of senescence, as determined based on the expressed RNAs. The review of Li et al.[4] discusses use of signatures to detect organ- and cell type-specific senescence, since senescent cells exhibit marked heterogeneity across tissues due to differences in developmental origin, cell composition, extracellular milieu and exposure to age- or stress-induced damage. Investigating cellular senescence signatures, helps in the understanding of organ-specific contexts due to the diverse biological functions and cellular compositions of different tissues.

Signatures are also the basis for molecular profiling and other forms of precision and personalized medicine, e.g., the Oncotype DX[5], Foundation Medicine[6] and IMPACT signatures[7] in cancer.

Finally in the domain of aging science Kraus et al[8] present a small RNA signature and models that predict 2 year survival in older healthy adults and have strong estimated effects on survival making them suitable for new therapeutics R&D.

**Revisiting and Comparison to Venet et al Study Findings**

The Venet et al study[9] findings which were focused on breast cancer bulk transcriptomics and a reference proliferation signature, at face value suggest that (1) random signatures correlate with the phenotype and (2) the proportion of published signatures that are matched or outperformed in terms of p-value by random marker selection is 60%, much higher than most of the reported proportions for most of the feature section methods and estimator combo in our analyses (0%-79%, with median of 14% depending on estimator and feature selection method used, Table 1).

With the benefit of our theoretical framework we can verify the former claim and reject the latter discrepancy with our empirical results as not caused by a special characteristic of breast cancer bulk transcriptomics versus the wide range of omics types and phenotypes examined in the present report. Rather, the Venet et al 60% estimate is artefactual due to a combination of systematic and random biases in how the random testing was conducted, that upward-biased (inflated) the reported performance of random gene selection.

The following biases can be observed:

(a) Systematic bias #1: Non-random (i.e., previously published) signatures were not tailored to the population from which the validation set is sampled (but to a different sample from a different population). This is an example of underfitting/under performant models bias due to discovery-application target population mismatch[10]. This mismatch systematically lowers the signal of the non random signatures. Per our analysis, the lower the predictivity of algorithmic or expert signatures, the easier it is for a random signature to match and exceed them. That is why our empirical experiments focused strictly on optimal signature predictivity comparisons on the same datasets.

(b) Systematic bias #2: models were not fitted to optimize predictivity but flowed a convenience protocol based on first Principal Component dimensionality reduction and Cox Regression,

(c) Systematic bias #3: Proliferation. Venet et al did not test their proliferation signature against other functional annotations of either algorithmic or random signatures that perform well. Our empirical analysis of the MB equivalence classes in real data shows that each member of the optimal signature equivalence class (which may be picked up by either algorithms, expert assessment or random selection) relates to one or more major functional categories. Thus one can create reference signatures for any functional category (proliferation or otherwise) and confirm that the signature performs well. Such a result is non-specific however (i.e,., proliferation or other functional annotation does not have special status).

(d) Random bias: Single-dataset evaluation. Venet et al testing of the p-value delta between random and published non-random signatures was conducted on a single dataset which cannot possibly be representative of the whole spectrum of bulk transcriptomics populations and datasets sampled from them for breast cancer and certainly not outside breast cancer and other assay types.

In conclusion, while the main thesis of the Venet paper is correct, the specific estimates about random signature p-values were upward biased due to a combination of systematic and random biases in their analyses.

1. Casella, G. *et al.* Transcriptome signature of cellular senescence. *Nucleic Acids Res.* **47**, 7294–7305 (2019).
2. Gorgoulis, V. *et al.* Cellular Senescence: Defining a Path Forward. *Cell* **179**, 813–827 (2019).

3. Wechter, N. *et al.* Single-cell transcriptomic analysis uncovers diverse and dynamic senescent cell populations. *Aging* **15**, 2824–2851 (2023).
4. Li, S. *et al.* Advancing biological understanding of cellular senescence with computational multiomics. *Nat. Genet.* **57**, 2381–2394 (2025).
5. Carlson, J. J. & Roth, J. A. The impact of the Oncotype Dx breast cancer assay in clinical practice: a systematic review and meta-analysis. *Breast Cancer Res. Treat.* **141**, 13–22 (2013).
6. Milbury, C. A. *et al.* Clinical and analytical validation of FoundationOne®CDx, a comprehensive genomic profiling assay for solid tumors. *PloS One* **17**, e0264138 (2022).
7. Cheng, D. T. *et al.* Memorial Sloan Kettering-Integrated Mutation Profiling of Actionable Cancer Targets (MSK-IMPACT): A Hybridization Capture-Based Next-Generation Sequencing Clinical Assay for Solid Tumor Molecular Oncology. *J. Mol. Diagn. JMD* **17**, 251–264 (2015).
8. Kraus, V. B. *et al.* Select Small Non-Coding RNAs Are Determinants of Survival in Older Adults. *Aging Cell* **25**, e70403 (2026).
9. Venet, D., Dumont, J. E. & Detours, V. Most Random Gene Expression Signatures Are Significantly Associated with Breast Cancer Outcome. *PLOS Comput. Biol.* **7**, e1002240 (2011).
10. Aliferis, C. & Simon, G. Overfitting, Underfitting and General Model Overconfidence and Under-Performance Pitfalls and Best Practices in Machine Learning and AI. in *Artificial Intelligence and Machine Learning in Health Care and Medical Sciences: Best Practices and Pitfalls* (eds Simon, G. J. & Aliferis, C.) (Springer, Cham (CH), 2024).
